# nucMACC: A MNase-seq pipeline to identify structurally altered nucleosomes in the genome

**DOI:** 10.1101/2022.12.29.521985

**Authors:** Wernig-Zorc Sara, Kugler Fabian, Schmutterer Leo, Räß Patrick, Hausmann Clemens, Holzinger Simon, Längst Gernot, Schwartz Uwe

**Affiliations:** Regensburg Center for Biochemistry (RCB), University of Regensburg, Regensburg, Germany; St. Anna Children’s Cancer Research Institute (CCRI), Vienna, Austria; NGS Analysis Center Biology and Pre-clinical Medicine, University of Regensburg, Regensburg, Germany

## Abstract

Micrococcal nuclease sequencing (MNase-seq) is the state-of-the-art method for determining chromatin structure and nucleosome positioning. Data analysis is complex due to the AT-dependent sequence bias of the endonuclease, and the requirement for high sequencing depth. Here, we present the newly developed nucleosome-based MNase accessibility (nucMACC) pipeline unveiling the regulatory chromatin landscape by measuring nucleosome accessibility and stability. nucMACC represents the first systematic, and genome-wide approach for detecting unstable (“fragile”) nucleosomes.

We characterized the regulatory nucleosomal landscape in *D. melanogaster* and *S. cerevisiae*. Two functionally distinct sets of promoters were identified, one associated with an unstable nucleosome and the other being nucleosome depleted. Chromatin structure analysis shows that unstable nucleosomes present intermediate states of nucleosome remodeling, preparing inducible genes for transcriptional activation in response to stimuli or stress. The presence of unstable nucleosomes correlates with RNA polymerase II proximal pausing. The nucMACC pipeline offers unparalleled precision and depth in nucleosome research and is a valuable tool for future nucleosome studies.

**Teaser:** The nucMACC pipeline quantifies the local and global functional alterations of chromatin structure.

## INTRODUCTION

Nucleosomes are the basic unit of DNA packaging in eukaryotic cells. First electron microscopy images of chromatin spreads displayed regularly spaced nucleosome core particles, separated by a short linker DNA, appearing like “beads-on-a-string” (Olins *et al*, 1975). The nucleosome core particle consists of 147 bp of DNA wrapped around a histone octamer, composed of the histone proteins H2A, H2B, H3, and H4 (Luger *et al*, 1997; Kornberg, 1974). The stability of the nucleosome core is maintained by over 360 hydrogen bonds between the DNA and histone proteins (Davey *et al*, 2002). Linker DNA varies in length from 15 to 95 bp depending on the species, cell type, and even the chromatin state within the same nucleus (Valouev *et al*, 2011; Compton *et al*, 1976; Prunell & Kornberg, 1982). DNA in the linker is generally accessible to DNA-binding factors, whereas access to DNA wrapped around the histone octamer is hindered. Nucleosome positioning, structure, and modification play a critical role in regulating gene expression, determining the accessibility of DNA to DNA-binding factors, thereby defining cell type-specific gene activity and repression (Valouev *et al*, 2011; Teif *et al*, 2012). *In vitro* experiments have shown that nucleosome positioning depends on the bendability and structure of the underlying DNA sequence and the effect of steric hindrance between neighboring nucleosomes (Struhl & Segal, 2013; Kaplan *et al*, 2009). On the other hand, *in vivo* nucleosome positioning is dynamically modified and influenced by additional factors, such as chromatin remodelers, histone variants, transcription factors (TF), histone post-translational modifications, or the transcriptional machinery (Chereji & Clark, 2018). Regulatory regions, such as promoters, enhancers, and insulators, are characterized by well-positioned nucleosomes, which is a common feature of eukaryotic chromatin (Hu *et al*, 2011; Schones *et al*, 2008).

In addition to the positioning of nucleosomes, the structural property of the nucleosomes is central to DNA accessibility and chromatin function. Nucleosomes can vary in their histone composition and number. Histone variants modify the structural and functional properties of nucleosomes, playing specific roles in chromatin regulation, and being incorporated into nucleosomes independent of the cell cycle (Talbert & Henikoff, 2017). For example, the histone H3 variant CENP-A is incorporated into nucleosomes at the centromeres. DNA-histone interactions are weakened in CENP-A-containing nucleosomes, partially unwinding DNA from the histone surface (Hasson *et al*, 2013). In addition, the human testis-specific histone H3 variants, H3T and H3.5, form nucleosomes with reduced stability, as compared to nucleosomes with the canonical H3, as does the C-terminal half of H2A.Z1 (Sato *et al*, 2020; Urahama *et al*, 2016; Tachiwana *et al*, 2010). Hexasomes lacking an H2A-H2B dimer, can be formed by a variety of mechanisms including transcription, histone assembly, and chromatin remodeling (Kireeva *et al*, 2002; Prasad *et al*, 2016; Liu *et al*, 2019; Ramachandran *et al*, 2017; Hsieh *et al*, 2022). The preferential presence of hexasomes near transcription start sites suggests a potential role in gene regulation (Rhee *et al*, 2014). However, it is still unclear whether these altered/sub-nucleosomal structures are stable or short-lived nucleosome intermediates *in vivo*. Furthermore, there are several studies revealing structurally altered nucleosomes, like lexosome, alterosome, pre-nucleosome, tetrasome and reversosome, that could play specific roles in maintaining an active genome (Lavelle & Prunell, 2007; Fei *et al*, 2015; Ramachandran & Henikoff, 2016).

MNase-seq has been widely used to study chromatin structure and nucleosome positioning. It involves the use of micrococcal nuclease (MNase), an enzyme that selectively cleaves linker DNA, leaving the nucleosome core particles intact. The resulting DNA fragments are sequenced and analyzed to determine the location and distribution of nucleosomes along the genome (Baldi *et al*, 2020). MNase-seq provides a high-resolution map of the nucleosomal landscape and has contributed to our understanding of the dynamic and regulated nature of chromatin structure (Valouev *et al*, 2011; Teif *et al*, 2012; Diermeier *et al*, 2014). Initially, high MNase digestion conditions were used to completely hydrolyze chromatin to the mono-nucleosomal level. Later, we and others combined different MNase digestion conditions to study chromatin accessibility and nucleosome stability (Schwartz *et al*, 2018; Henikoff *et al*, 2011; Mieczkowski *et al*, 2016; Chereji *et al*, 2019, 2015; Schwartz *et al*, 2023). Due to the complexity of data analysis in MNase titration experiments, new metrics have been introduced to provide a stable framework for accurate quantification of chromatin accessibility and nucleosome occupancy (Mieczkowski *et al*, 2016; Chereji *et al*, 2019). However, up to six different MNase concentrations, from very low to high, were used per sample and additional spike-ins were suggested to be necessary, resulting in a high workload, high costs, and requiring large amounts of sample material. Furthermore, the lack of an established analytical pipeline makes it difficult to adapt these kinds of experiments to new sample formats.

Here, we present a comprehensive and user-friendly nucleosome-based MNase accessibility (nucMACC) analysis pipeline. The nucMACC workflow starts directly with raw sequencing data, assesses the critical quality metrics relevant to MNase-seq experiments, and summarizes the results into user-friendly reports (Figure S1). The workflow reveals the genome-wide nucleosome positions and their corresponding nucleosome accessibility (nucMACC) scores. The pipeline identifies hyper-and hypo-accessible nucleosome positions with exceptionally high or low nucMACC scores. In addition, as a novel feature of the nucMACC pipeline, nucleosome stability scores are inferred from sub-nucleosomal fragments identifying unstable or non-canonical nucleosomes. We show that the pipeline achieves robust results using only two MNase conditions per sample. In summary, the nucMACC pipeline is written in nextflow, available on GitHub (https://github.com/uschwartz/nucMACC), and integrates Docker software container, making it easy to use, portable, reproducible, and scalable.

## RESULTS

The nucMACC pipeline requires a minimum of two MNase-seq titration conditions as input, preferentially in combination with a histone immunoprecipitation step, to exclude non-histone protein-mediated DNA-binding events. MNase, which preferentially hydrolyzes accessible linker DNA between nucleosomes, releases nucleosome core particles from the chromatin. Depending on the MNase digestion conditions, nucleosomes are released at different rates (Figure 1A). At mild MNase conditions, nucleosomes from accessible and active genomic sites are preferentially released, referred to as hyper-accessible nucleosomes (Figure 1A, green nucleosome) (Schwartz *et al*, 2018; Mieczkowski *et al*, 2016). Increasing the MNase concentration or digestion time, increasingly releases nucleosome core particles from chromatin, until the nucleosomal chain is fully hydrolyzed. Furthermore, MNase is a processive enzyme. Once the nucleosome is released from chromatin, MNase progressively trims the linker DNA and then continues – at a lower rate – to hydrolyze the histone bound DNA, resulting in sub-nucleosomal fragments. Canonical nucleosome core particles are highly resistant to intra-nucleosomal MNase cleavage, and the nucleosome core DNA remains mainly complete, even at high MNase digestion conditions. However, non-canonical nucleosomes, often referred to as unstable or fragile nucleosomes, exhibit an increased MNase sensitivity and are degraded to sub-nucleosome-sized fragments already at low MNase concentrations (Figure 1A, blue nucleosome) (Kubik *et al*, 2015; Weiner *et al*, 2010; Voong *et al*, 2016). We refer to these kinds of nucleosomes as unstable nucleosomes in the remainder of this manuscript. At high MNase conditions, unstable nucleosomes are fully degraded and are therefore lost in this kind of MNase dataset. Hence, sub-nucleosomal fragments can be used to measure nucleosome stability in MNase titration experiments, as reflected by resistance to internal MNase cleavage.

**Figure 1.**
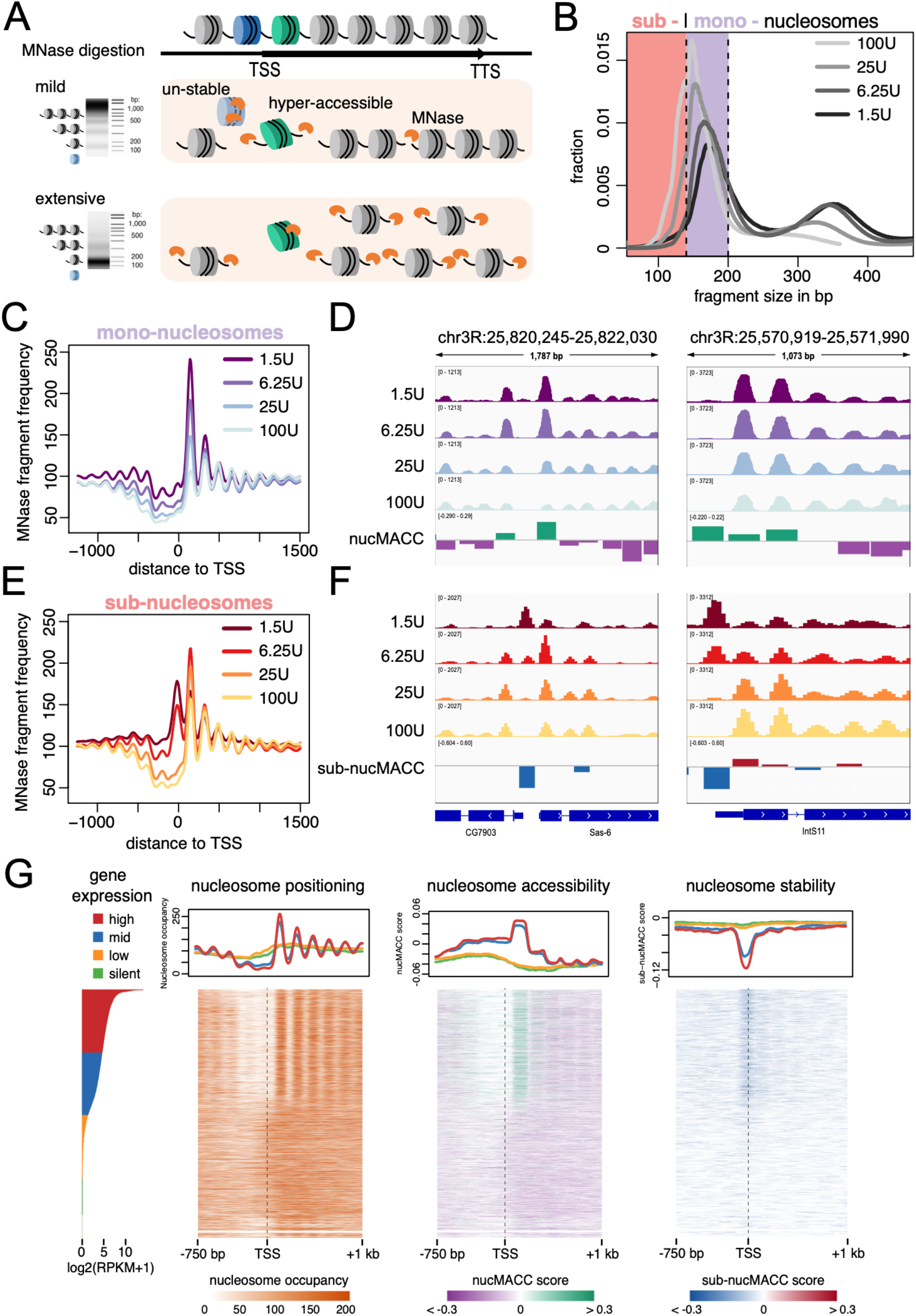
(A) Schematic representation of differential MNase chromatin digestion. Under mild MNase conditions, chromatin is partially digested and nucleosomes (green, hyper-accessible) from accessible regions are preferentially released (upper panel). At extensive MNase conditions, chromatin is completely digested to nucleosome core particles and MNase-sensitive nucleosomes (blue, unstable) are lost. (B) Fragment size distribution of MNase titrations. Fragments are selected based on the fragment size and grouped into sub-(< 140 bp, light red section) or mono-nucleosomes (140-200 bp, purple section). (C) and (E) average fragment frequencies at the TSS. (D) and (J) Genome browser snapshot showing the output of the nucMACC pipeline. nucMACC scores are deduced from mono-nucleosomal fragments (D) and sub-nucMACC scores from sub-nucleosomal fragments, respectively (J). (E) Heatmaps sorted by gene expression showing nucleosome positioning (left), accessibility (middle, nucMACC scores), and stability (right, sub-nucMACC scores) at TSS.

Starting from raw sequencing files in the FASTQ format, the nucMACC pipeline first assesses the sequencing quality. Next, the reads are aligned to the provided reference genome and subsequently mapped reads are filtered based on mapping quality (Figure S2). Optionally ambiguous genomic elements, such as blacklisted regions or mitochondrial chromosomes, can be removed from the analysis. To obtain high-resolution nucleosome positions, fragments between 140 and 200 bp in length, the typical size of mono-nucleosomal DNA, are selected (Figure 1B). Size-selected fragments of all MNase titration conditions are pooled and used to derive a comprehensive map of the genome-wide nucleosome positions.

Recently, a metric to measure chromatin accessibility, termed MACC (MNase accessibility), was introduced (Mieczkowski *et al*., 2016). As the MACC scores provide an elegant way to measure chromatin accessibility based on MNase-seq data, we adapted the MACC scoring system but changed the principle of calculation to obtain a higher annotation resolution and accuracy. Instead of counting the MNase reads in arbitrary genomic bins as in the original MACC version, we quantify the MNase accessibility directly at the defined nucleosome positions, which results in a specific accessibility score for each individual nucleosome (nucMACC). Next, linear regression is conducted on the normalized fragment frequencies for each nucleosome (Figure S2, see Methods section for details). The slope of the regression line multiplied by minus one is used as the nucMACC score. Since MNase exhibits an increasing AT preference at higher concentrations, raw nucMACC scores are normalized for the underlying GC content using a LOESS regression (Schwartz *et al*, 2018).

As a new feature of the nucMACC pipeline fragments shorter than 140 bp, representing sub-nucleosomal DNA fragments, are used to measure nucleosome stability. We use only the lowest MNase concentration to call these sub-nucleosomal positions (called sub-nucleosomes in the remaining text), as at higher MNase titration conditions, these unstable nucleosomes are fully hydrolyzed (Figure 1A, blue nucleosome). Analogous to the nucMACC scores, sub-nucleosomal fragment frequencies for each MNase titration condition are quantified at the defined sub-nucleosome positions. The slope of the regression is used to measure nucleosome stability and normalized to the underlying GC content. The normalized score is referred to as the sub-nucleosome MNase accessibility score (sub-nucMACC).

### The nucMACC pipeline scores nucleosome accessibility and stability

To establish and validate the nucMACC pipeline, we used the MNase chromatin digestions coupled with H4 immunoprecipitation from the original MACC publication (Mieczkowski *et al*, 2016). The dataset collection consists of four different MNase concentration conditions (1.5U, 6.25U, 25U, and 100U), performed in two replicates each. The fragment size distribution is an important quality control parameter, as it clearly shows the degree of chromatin hydrolysis in each sample (Figure 1B). With increasing MNase concentrations the di-nucleosome fraction is disappearing, and the levels of mono-nucleosomes are increasing. The dominant mono-nucleosomal peak is in the range of 140–200 bp, while the peak summit shifts with higher MNase concentrations closer to 147 bp, the protected DNA-length of a nucleosome core particle. Fragment sizes shorter than 140 bp indicate sub-nucleosomal DNA fragments, which mainly arise from DNA cleavage within the nucleosomal core. Sub-nucleosomal DNA fragments are barely present at mild MNase conditions but increase with higher MNase concentrations. Based on the shape of the fragment size distribution curve, we selected the fragments in the range of 140 – 200 bp in length for mono-nucleosome analysis and fragments shorter than 140 bp for sub-nucleosome analysis, respectively (Figure 1B).

Plotting the fragment frequencies of mono- and sub-nucleosomes at the TSS shows that the selected categories behave differently, suggesting different structural information or functional behavior (Figure 1C and 1E). Analyzing the mono-nucleosomes of the two high MNase conditions (25U and 100U) reveals the well-known nucleosome positioning pattern at the TSS: a nucleosome depleted region (NDR) followed downstream by a phased nucleosome array, which starts with a well-positioned +1-nucleosome and spreads into the gene. The low MNase conditions (1.5U and 6.25U) reveal the same mono-nucleosomal positions, however, specific positions, such as the +1-nucleosome, are markedly enriched compared to the high MNase conditions. This effect clearly indicates that nucleosomes, depending on their genomic location, exhibit different release rates from chromatin. The nucMACC score provides a quantitative assessment of chromatin accessibility, revealing the change in mono-nucleosome fragment frequencies with increasing MNase concentration (Figure 1D and S2). Comparing the mono-nucleosome and the sub-nucleosome TSS profile at high MNase conditions, reveals a very similar pattern, suggesting that once the nucleosomes were released from chromatin, MNase trims the nucleosomal DNA. However, in the sub-nucleosomal fraction, an additional dominant peak at the denominated NDR appears at low MNase conditions. As the analyzed dataset is based on a H4-ChIP, these observations suggest that the NDR is not free of histones but occupied by MNase sensitive nucleosomes (Figure 1E). The sub-nucMACC score allows the quantification of the nucleosome stabilities at these sites, as it monitors the changes in sub-nucleosome fragment frequencies over MNase concentrations (Figure 1F and S2).

Even though the nucMACC and sub-nucMACC scores are calculated in a similar manner, their scores do not correlate at overlapping positions, confirming that both metrics reveal distinct nucleosomal features (Figure S3A). The correlation of the nucMACC scores with the original MACC scores (Mieczkowski *et al*, 2016) was modest (R=0.48), which can be explained by the fact that the scores in our analysis are centered on the actual nucleosome positions revealing functional details with high resolution. The sub-nucMACC scores, which is a novel category implemented in our pipeline, showed no correlation (R=-0.24) indicating new features of the dynamic chromatin structure (Figure S3B-C).

By centering the nucMACC scores on the TSS, we show that the +1 nucleosomes of actively transcribed genes are in general hyper-accessible, whereas nucleosomes further downstream in the gene body did not significantly change nucleosome accessibility (Figure 1G). Remarkably, in genic regions, we observed no difference in accessibility between silent and highly expressed genes (Figure S3D), confirming that chromatin accessibility is not changed at a global scale but modulated only locally, changing the accessibility of individual nucleosomes at regulatory sites, such as the promoter (Schwartz *et al*, 2018).

The nucleosome stability profile at the TSS of active genes was different compared to the nucleosome accessibility score. The lowest nucleosome stability scores were obtained upstream of the +1 nucleosome directly at the TSS of active genes. The sub-nucleosome analysis uncovered unstable nucleosomes directly at the NDR, showing an overall low nucleosome occupancy in mono-nucleosome analysis (Figure 1G). In contrast, the transcription end sites (TES) are nucleosome depleted in *Drosophila*, as no unstable nucleosomes can be detected (Figure S3E), in contrast to studies performed in yeast and mouse (Voong *et al*, 2016; Xi *et al*, 2011).

Overall, the nucleosome features provided by the nucMACC pipeline are robust and reproducible, as shown by the comparison to other MNase titration experiments that were combined with a H3-ChIP (Figure S4A-G) (Mieczkowski *et al*, 2016). We observed the highest variation between the H3- and the H4-MNase-ChIP experiments in the number and location of the sub-nucleosome positions (Figure S4C). This can be explained, by the significantly lower number of sub-nucleosomal fragments in the H3 low MNase samples compared to the H4 dataset, giving rise to fewer called sub-nucMACC positions (addressed in more detail below). Nevertheless, the sub-nucMACC scores at overlapping positions revealed a high correlation (Figure S4C, R=0.8).

In summary, splitting up MNase titration data by fragment sizes into mono-nucleosomes and sub-nucleosomes enables the nucMACC pipeline to reveal nucleosome positions, nucleosome accessibility, and nucleosome stability scores within the same experiment.

Next, we tested whether the additional histone immunoprecipitation step after chromatin digestion is required to obtain reliable nucleosome features. Without ChIP, mono- and sub-nucleosome positioning changed and overlapping positions showed only a modest correlation with whole chromatin extractions (Fig S4H-K). Changes in the fragment profile were clearly visible upstream of the TSS, and most pronounced at the -1 nucleosome position affecting nucMACC and sub-nucMACC scores (Figure S4L-N). The -1 nucleosome position exhibited the highest accessibility throughout all conditions, whereas the sub-nucleosomal fraction exhibited a broad and pronounced enrichment at the promoter of expressed genes in the low MNase concentrations. This result is reminiscent to other methods measuring chromatin accessibility, such as DNase- or ATAC-seq (Vierstra *et al*, 2014; Buenrostro *et al*, 2013). The differences between whole chromatin analysis and the combined MNase-ChIP datasets can be explained by non-histone proteins bound to DNA (Mieczkowski *et al*, 2016). Our analysis shows that an additional histone immunoprecipitation step is required, to obtain accessibility and stability measurements specifically relating to nucleosomes. Nevertheless, the nucMACC pipeline can be run using whole chromatin extractions and measures overall chromatin accessibility and stability over all molecules bound to DNA.

### nucMACC scores identify hyper- and hypo-accessible nucleosomes

To identify functionally important nucleosomes, the nucMACC pipeline extracts nucleosomes with exceptionally high or low accessibility, termed hyper- or hypo-accessible nucleosomes. For this, the nucMACC score of each nucleosome is plotted against its rank. Ranks are divided by the total number of nucleosomes and the nucMACC score, by the total range of scores. In that way, both the x and y axes have a value range of 1 (Figure 2A, Supplemental Dataset 1). To geometrically identify the positions, where the signal decreases/increases rapidly, we determined the positions at the curve where the slope is the first/last time below one. Hypo-accessible nucleosomes are defined as the nucleosome positions before the slope of the curve decreases below 1 (Figure 2A, purple dots), and hyper-accessible nucleosomes after the slope increases again above 1, respectively (Figure 2A, green dots). A similar strategy was previously applied to define super-enhancers (Whyte *et al*, 2013).

**Figure 2.**
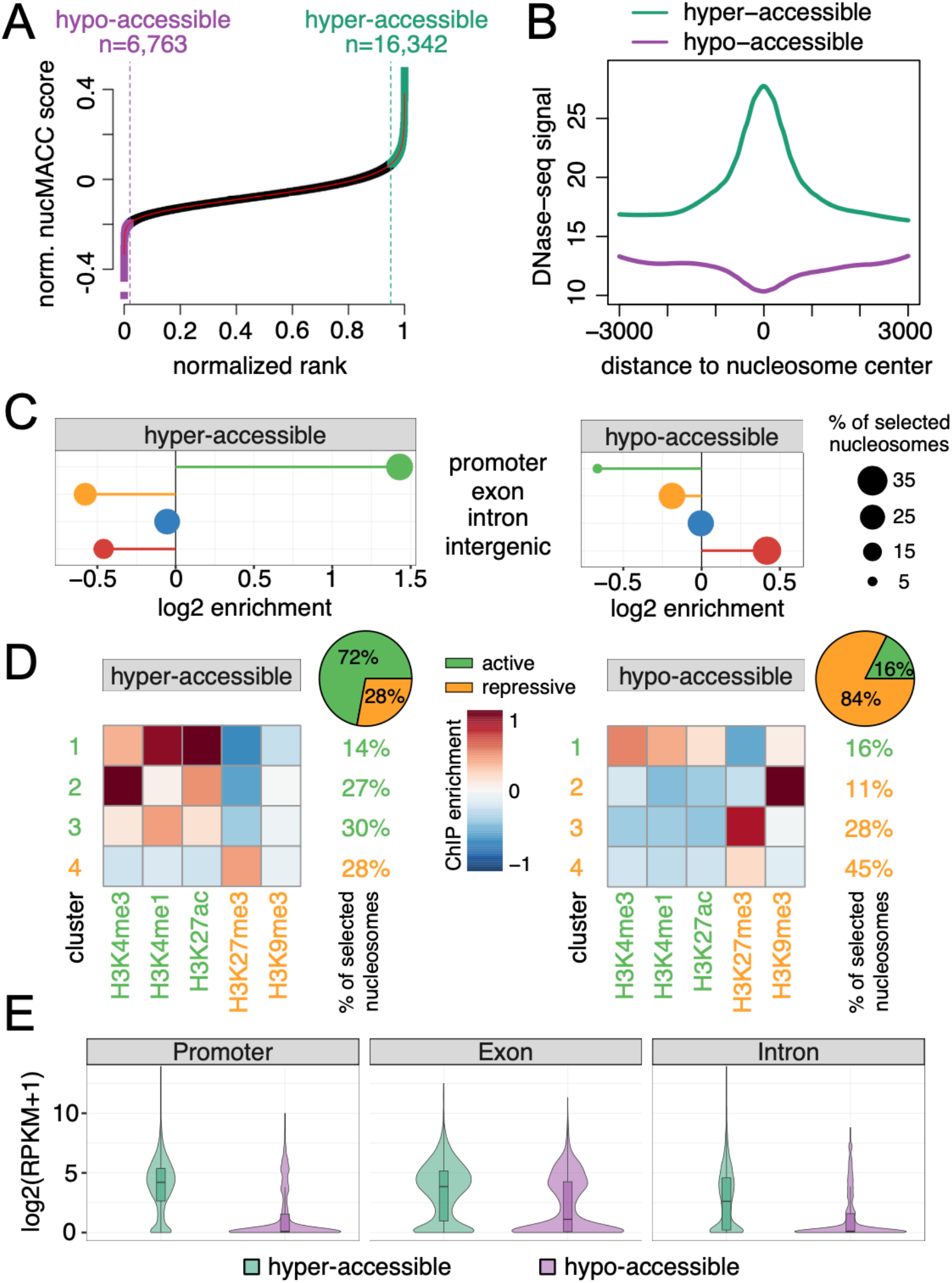
(A) Selection of hyper-(green) and hypo-accessible (purple) nucleosomes based on nucMACC scores. Nucleosomes were ranked by nucMACC score. X-and y-axis were normalized to a range of one. Cut-offs were deduced from the slope of the LOESS curve fit. Nucleosomes before the point, where the slope goes the first time below 1 (purple dashed line) were classified as hypo-accessible. Nucleosomes after the point where the slope of the curve exceeds one again (green dashed line), were classified as hyper-accessible nucleosomes. (B) Correlation of DNase-seq sites with hyper-/hypo-accessible nucleosomes. (C) Enrichment of hyper-/hypo-accessible nucleosomes at genomic features. (D) K-means clustering (n=4) of histone modifications at hyper-/hypo-accessible sites. Centroid values of each cluster are indicated by the color code. The relative number of nucleosomes represented in each cluster is indicated on the right side. Repressive histone modifications are highlighted in orange and active in green, respectively. The pie chart depicts the fraction of nucleosomes assigned to either active or repressive histone clusters. (E) Gene expression levels of genes exhibiting hyper-/hypo-accessible nucleosomes at distinct features. Promoter was defined as 500 bp upstream of the TSS to 300 bp downstream.

In total, we identified 6,763 (2 %) hypo-accessible and 16,342 (4,9 %) hyper-accessible nucleosomes in the *Drosophila* S2 H4-MNase-ChIP data set. Hyper-accessible nucleosomes are strongly enriched at DNase I hypersensitive sites, whereas hypo-accessible nucleosomes are depleted, confirming the nucMACC score as a measure for general DNA accessibility (Figure 2B). Genomic feature characterization of hyper- and hypo-accessible nucleosomes revealed a significant enrichment at promoter and intergenic regions, respectively (Figure 2C). Hyper-accessible nucleosomes are preferentially associated with H3K4me1, H3K4me3, and H3K27ac histone modifications, marking active TSS and enhancer, whereas hypo-accessible nucleosomes are associated with H3K9me3 and H3K27me3 modifications, indicating heterochromatic regions (Figure 2D). In agreement with the histone modification analysis, hyper-accessible nucleosomes occupy mainly expressed genes, while hypo-accessible nucleosomes are present on silent genes (Figure 2E). Nevertheless, we identified a substantial fraction of hypo-accessible nucleosomes at exons of expressed genes, potentially regulating differential exon usage.

The nucMACC pipeline identifies functional nucleosome positions at regulatory sites of the genome, showing that the regulation of genome activity is associated with differential nucleosome accessibility.

### sub-nucMACC scores identify the presence of non-canonical and unstable nucleosomes

To measure nucleosome stability, sub-nucleosome positions were derived from the sub-nucleosome fragments in the lowest MNase concentration condition. Hyper-accessible nucleosomes are expected to have a higher MNase degradation rate as they are less protected once they are released from chromatin. Here, sub-nucleosome positions are selected, which either contained at least four-fold more normalized sub-nucleosome fragment counts than the mono-nucleosome fraction or did not overlap with any mono-nucleosome position (Figure S1).

Like the selection of hyper- and hypo-accessible nucleosomes, nucleosomes exhibiting extreme stability scores (sub-nucMACC) were selected (Figure 3A, Supplemental Dataset 1). This selection strategy resulted in 4,233 (6.7 %) nucleosomes showing very low stability scores and therefore termed unstable nucleosomes. Additionally, a rather small subset of nucleosomes (n=355, 0.6 %) were enriched in the sub-nucleosome fraction of the lowest MNase condition and remained stable at higher MNase concentrations. This group represents most likely structurally altered nucleosomes with additional MNase cleavage sites in the realm of the nucleosome core or non-octamer histone compositions, such as hexa- or hemisomes (Hasson *et al*, 2013; Rhee *et al*, 2014; Dalal *et al*, 2007). Nucleosomes enriched in the low sub-nucleosomal fractions and showing extraordinarily high stability scores are referred here to as non-canonical nucleosomes.

**Figure 3.**
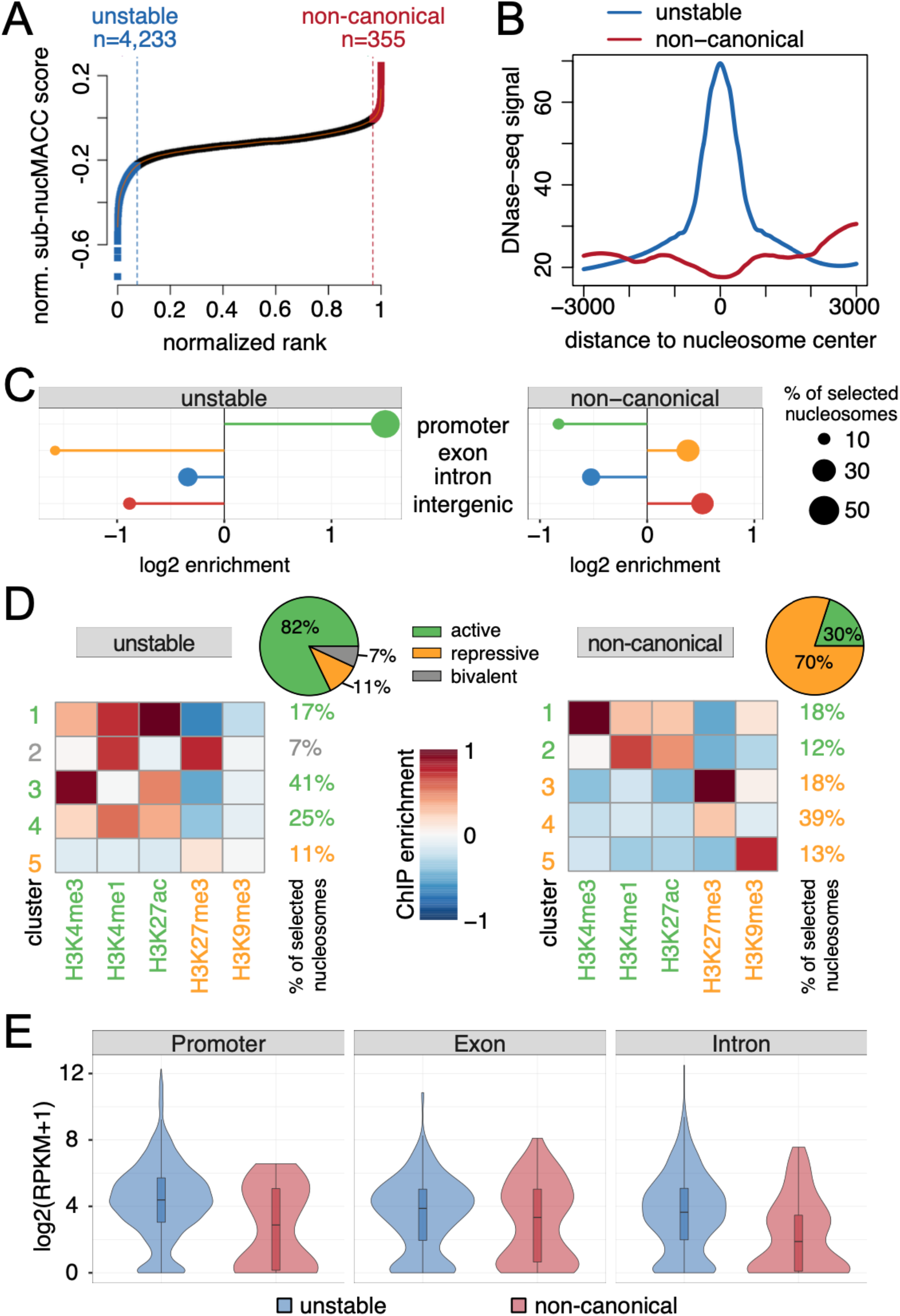
(A) Selection of unstable (blue) and non-canonical (red) nucleosomes based on nucMACC scores. Nucleosomes were ranked by sub-nucMACC score. X- and y-axis were normalized to a range of one. Cut-offs were deduced from the slope of the LOESS curve fit. Nucleosomes before the point, where the slope goes the first time below 1 (blue dashed line) were classified as unstable. Nucleosomes after the point where the slope of the curve exceeds one again (red dashed line), were classified as non-canonical nucleosomes. (B) Correlation of DNase-seq sites with unstable and non-canonical nucleosomes. (C) Enrichment of unstable or non-canonical nucleosomes at genomic features. (D) K-means clustering (n=5) of histone modifications at unstable or non-canonical sites. Centroid values of each cluster are indicated by the color code. The relative number of nucleosomes represented in each cluster is indicated on the right side. Repressive histone modifications are highlighted in orange, active in green, respectively. The bivalent cluster enriched for both repressive and active histone modifications is indicated in grey. The pie chart depicts the fraction of nucleosomes assigned to either active, repressive, or bivalent histone clusters. (E) Gene expression levels of genes exhibiting unstable or non-canonical nucleosomes at distinct features. Promoter was defined as 500 bp upstream of the TSS to 300 bp downstream.

In contrast to non-canonical nucleosomes, unstable nucleosome positions are characterized by a high chromatin accessibility (Figure 3B), being mainly located at gene promoters (53,8 %), whereas non-canonical nucleosomes are enriched in intergenic and exonic regions (Figure 3C). Preferentially, active histone marks are associated with unstable nucleosomes (Figure 3D), and accordingly, unstable nucleosomes are present at expressed genes (Figure 3E). Interestingly, we identify a subgroup of unstable nucleosomes associated with active H3K4me1 and repressive H3K27me3 marking bivalent enhancers. Conversely, non-canonical nucleosomes showed enrichment in repressive H3K27me3 and H3K4me1 (Figure 3D). Nevertheless, 30% of the non-canonical nucleosomes are associated with active histone modifications, and therefore non-canonical nucleosomes are located at both expressed and silent genes (Figure 3E).

### Unstable nucleosomes occupy promoters with high RNA polymerase pausing rate

As we observed that unstable nucleosomes are enriched upstream of the TSS (Figure 1G and Figure 3C) of expressed genes, we addressed their role in gene regulation. Therefore, we divided all expressed genes into two groups: 1) those that contained an unstable nucleosome 200 bp upstream of the TSS (n=1,588), and 2) those that exhibited a nucleosome-depleted region (NDR) in the promoter (n=5,363) (Figure 4A, Supplemental Dataset 1). To discriminate between NDR or unstable nucleosome containing promoters, it is obligatory to use MNase treatments coupled with histone immunoprecipitations, as DNA fragments protected by non-histone proteins would dominate the promoter signal in whole chromatin MNase digestion, resulting in sub-nucleosomal fragments at all expressed promoters (Figure S5). Overall, the steady-state transcript abundance of the corresponding genes was similar in both groups and only slightly elevated for genes carrying an unstable nucleosome (Figure 4B, p-value = 2x10^-15^). GO enrichment analysis revealed that genes containing unstable nucleosomes in their promoter are stimulus or developmentally regulated (Figure 4C). Furthermore, unstable nucleosome promoters were enriched for *Drosophila* promoter motifs Unknown1-3 and M1BP motif, but depleted in Initiator, TATA-box, and Unkown4 motif (Figure 4D). As M1BP is a known factor to regulate RNA Pol II pausing depending on nucleosome positioning (Li & Gilmour, 2013), it is suggested that RNA Pol pausing provides an efficient way to induce gene transcription rapidly or synchronously, in response to developmental and environmental signals. We tested whether Pol II pausing differed between NDR or unstable nucleosome promoter using nascent RNA sequencing data (Fuda *et al*, 2015). In agreement with M1BP motif enrichment, unstable nucleosome promoters show a pronounced association with paused Pol II directly downstream of the TSS and higher pausing index rates compared to the NDR promoters, enabling a fast transition into productive transcription at responsive and developmental genes (Figure 4E-D).

**Figure 4.**
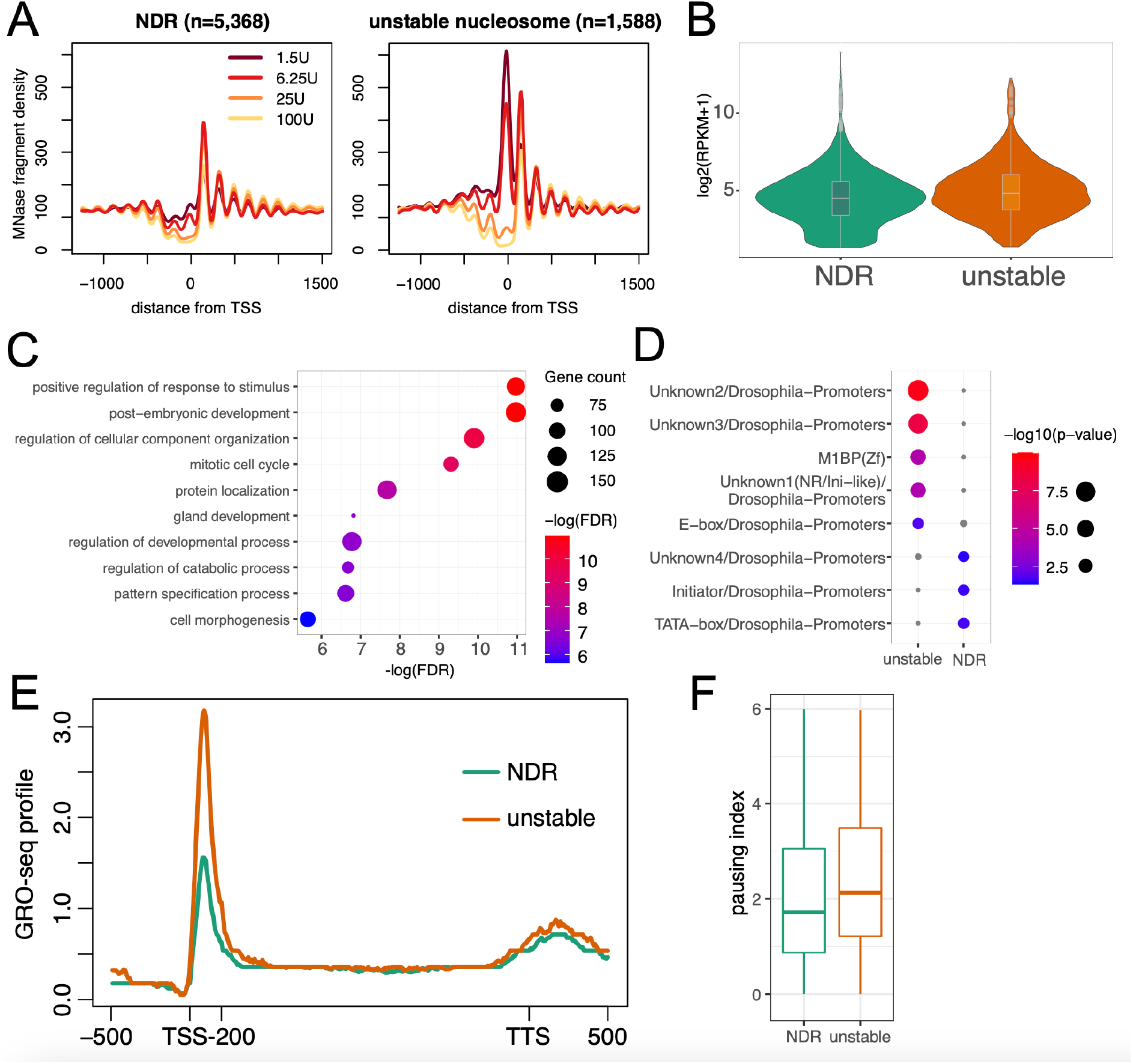
(A) Average fragment frequencies at the TSS of expressed genes either containing an unstable nucleosome (n=1,588, right) or exhibiting a nucleosome depleted region (NDR) (n=5,368, left) directly upstream of the TSS. (B) Violin plot showing expression levels of genes either containing an unstable nucleosome (n=1,588, right) or exhibiting a nucleosome depleted region (NDR) (n=5,368, left) directly upstream of the TSS. (C) Enrichment of biological processes associated with genes characterized by an unstable nucleosome promoter. (D) Motif enrichment analysis in the unstable nucleosome (left) or NDR (right) promoter. (E) Median GRO-seq profile showing nascent RNA abundance. Regions upstream of TSS, downstream of TTS and the first 200 bp downstream of TSS are unscaled. The remaining gene body is scaled to have the same length for each gene. (F) Boxplots showing the pausing index based on the GRO-seq signal. Pausing index was calculated as the fold change of the GRO-seq signal in the first 200 bp downstream of the TSS versus the remaining gene body.

### M1BP binds to DNA occupied by an unstable nucleosome

We next assessed the relationship between the binding of the transcription pausing factor M1BP and nucleosome properties. Here, we investigated nucleosome positioning, accessibility, and stability at annotated M1BP binding sites (Zouaz *et al*, 2017). We observed that M1BP binding sites are surrounded by hyper-accessible nucleosomes (Figure 5A-C). Directly in the binding center of M1BP, we identified a clear enrichment of unstable nucleosomes (Figure 5D-F). This observation suggests that M1BP does not bind to nucleosome-free DNA but rather to DNA, still co-occupied with histones like a pioneer transcription factor (Michael *et al*, 2020). Another possibility is that M1BP competes with nucleosomes at the same position and the signal is a mixture of two different populations of cells that have either a nucleosome or M1BP bound at that position. Regardless, the enrichment of unstable nucleosomes clearly shows that histone DNA interactions are apparently weakened making them sensitive to internal MNase digestions, which could be the result of transcription factor binding to the nucleosome core (Iwafuchi-doi & Zaret, 2014) or active remodeling (Brahma & Henikoff, 2019).

**Figure 5.**
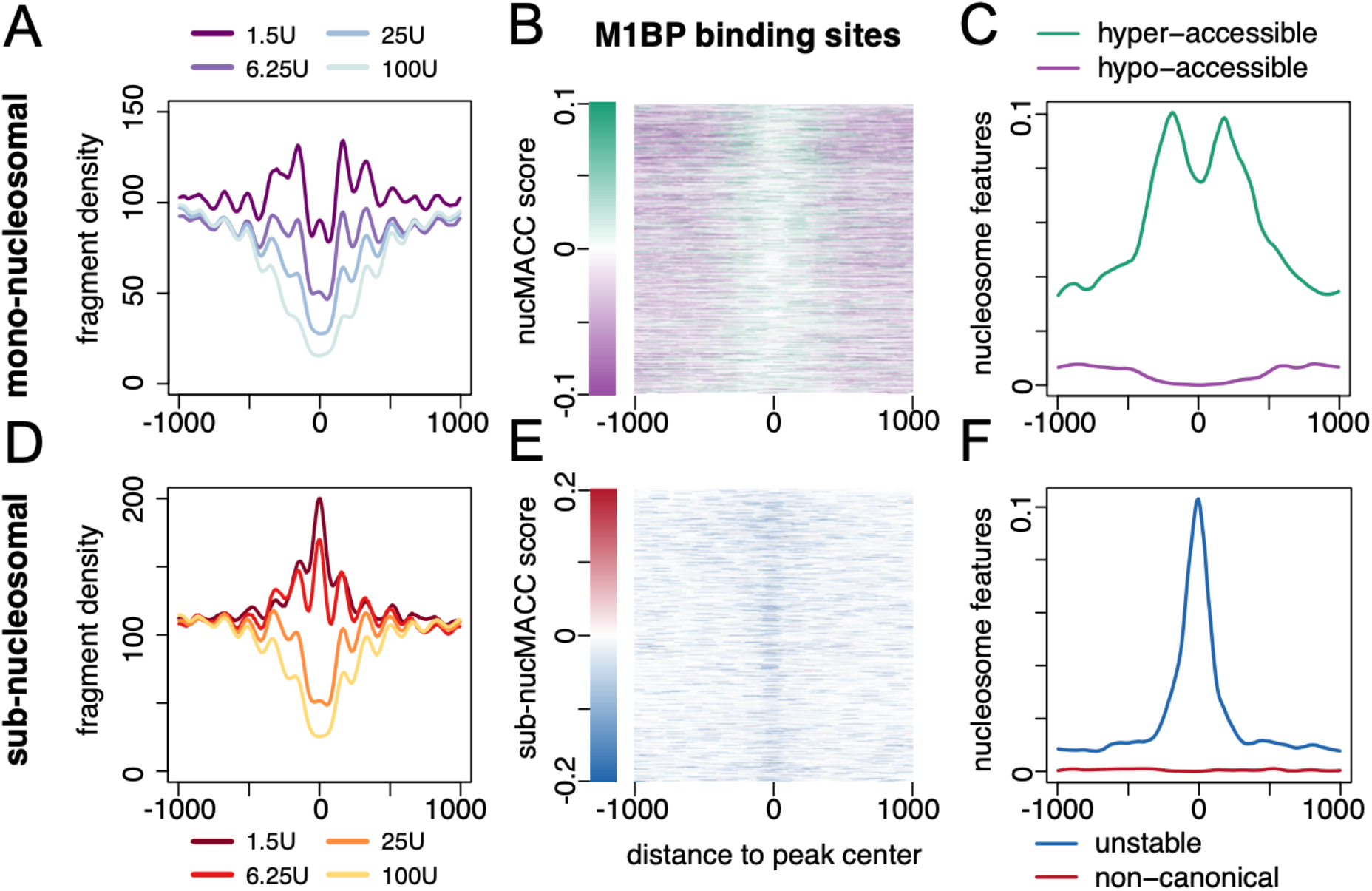
(A) and (D) average fragment frequencies of mono-(A) and sub-nucleosomal (D) fragments at M1BP bound sites. (B) and (E) Heatmap showing nucMACC (B) or sub-nucMACC (E) scores at M1BP bound sites. (C) and (F) Distribution of hyper-/hypo-accessible (C) or unstable and non-canonical (F) nucleosomes at M1BP bound sites.

### Evaluating the robustness of the nucMACC pipeline

Previously four or more MNase digestion conditions have been used to determine chromatin accessibility scores (Mieczkowski *et al*, 2016; Chereji *et al*, 2019). We sought to determine the minimum number of titration points per sample to obtain reliable and robust results with our pipeline. We compared the nucMACC and sub-nucMACC scores using two or four titration points of the same experimental dataset. As a result, we show that only two different MNase conditions are required, being highly correlated with the four titration conditions. This correlation holds true, if the difference in the compared MNase concentration conditions is at least 10-fold (Figure 6A and B; Figure S6A an B). In addition, we only observed consistent results, when using the lowest available MNase concentration to detect unstable nucleosomes within the sub-nucleosomal fragments. At higher concentrations, fragments of unstable nucleosomes are often lost due to their fragile nature.

**Figure 6.**
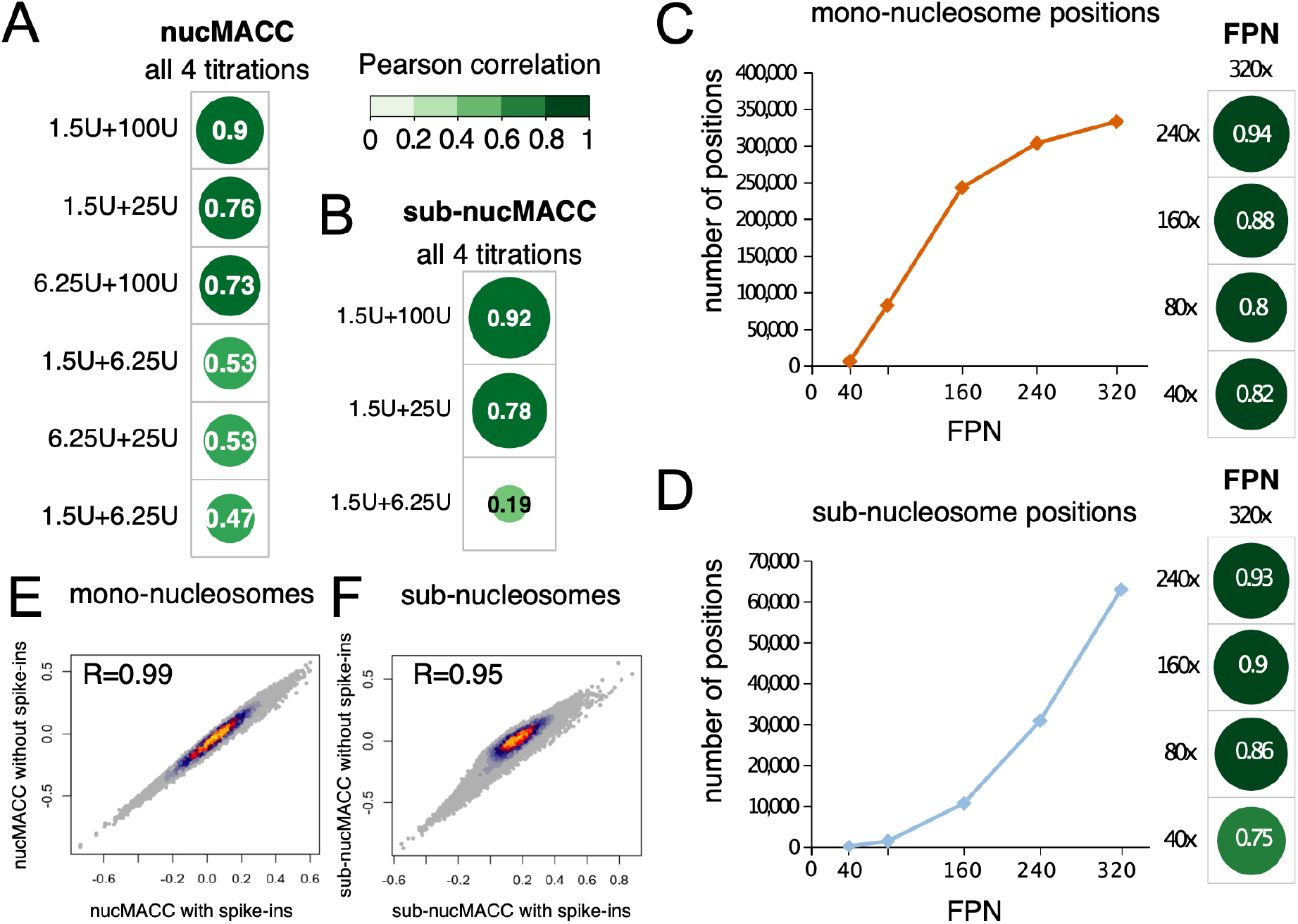
(A-B) Correlation of nucMACC (A) or sub-nucMACC (B) scores between samples with two or four MNase titrations. The pearson correlation coefficient is indicated (C-D) The number of called nucleosomes based on sequencing depth (FPN – fragments per nucleosome). Correlation of nucMACC (C) or sub-nucMACC (D) scores between samples with different sequencing depths is illustrated at the right. (E-F) Scatterdensityplot showing correlation of nucMACC (E) or sub-nucMACC (F) scores with or without spike-in information. The Pearson correlation coefficient is indicated.

Next, we addressed the importance of sequencing depth for the robustness of the nucMACC pipeline, as MNase-seq analysis usually requires a high sequence coverage of the genome. To obtain a genome independent quantification we normalized the sequencing depth to the number of fragments per nucleosome (FPN). To estimate the total number of nucleosomes we divided the size of the mappable genome by the expected nucleosome repeat length (here we used 170 bp as an approximation):

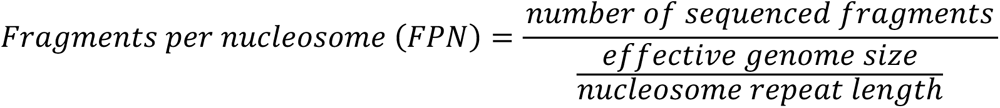

We found that using half of the fragments, with coverage of 160 FPN preserves 73% of the mono-nucleosome positions, and the nucMACC score correlation is reasonably high (R=0.88) (Figure 6C and Figure S6C). Whereas further lowering the sequencing depth to 80 FPN, results in a considerable loss of called mono-nucleosome positions to 25%, albeit the nucMACC score correlation remains stable at overlapping nucleosome positions, with a correlation of 0.80. For sub-nucleosomes, a value of 320 FPN results already at low coverage, since the sub-nucleosomal fragments are underrepresented in low MNase digestion conditions. Reducing the coverage to 240 FPN decreases the called sub-nucleosomes to 49% (Figure 6D). Nonetheless, the correlation between sub-nucMACC scores remains very high at overlapping positions (Figure S6D). Below 240 FPN coverage, the sub-nucleosome analysis produces an insufficient amount of called sub-nucleosome positions, not allowing meaningful analysis (Figure 6D). In summary, we recommend a minimum sequencing depth of 240 FPN or higher to be used when analyzing nucleosome stability.

Ultimately, we assessed the reliability and robustness of nucMACC and sub-nucMACC scores, testing if the differential amount of nucleosomes released at various MNase titration conditions would bias the scores. Therefore, we ran our pipeline on a published dataset with spiked-in nucleosomes and compared the results with or without spike-in information. Remarkably, we find almost no difference in nucMACC scores when comparing the pipeline with or without spike-in information for mono-(Figure 6E) and sub-nucleosomes (Figure 6F). In conclusion, the nucMACC pipeline does not require spike-ins and is robust and reproducible with only two MNase titrations.

### Unstable nucleosomes are associated with nucleosome remodeling

As the nucMACC pipeline produces consistent results with only two MNase titrations, we used appropiate published datasets to gain further insights into chromatin biology. A recent MNase-seq study in *Saccharomyces cerevisiae* questioned the existence of MNase-sensitive nucleosomes at promoter regions (Chereji *et al*, 2017). We used our nucMACC pipeline to re-analyze the H4-MNase-ChIP-seq data set starting from their raw sequencing files.

Investigating the results of the mono-nucleosome analysis, we observed that both hyper- and hypo-accessible nucleosomes are enriched at the promoter and predominantly at the +1 nucleosome position (Figure S7A, Supplemental Dataset 2). We selected the genes either associated with a hyper- or hypo-accessible +1 nucleosome and compared the initiation frequency and pausing rate using NET-seq data (Churchman & Weissman, 2011). In contrast to the results in *D. melanogaster*, genes associated with a hypo-accessible +1 nucleosome exhibited higher expression rates (Figure S7B). This can be explained by the rather homogenous chromatin accessibility in yeast, due to the high gene density and overall high transcriptional activity (Marr *et al*, 2021; Chen *et al*, 2016). Here, the local accessibility to MNase is mainly restricted by the local crowdedness of chromatin and additional non-histone proteins. Hypo-accessible +1 nucleosomes showed a higher occupancy of initiating RNA Pol II and factors of the initiation complex indicating increased DNA protection from MNase digestion by the RNA Pol II pre-initiation complex (Figure S7C). This is in agreement with the higher expression rates compared to hyper-accessible +1 nucleosomes. At normal conditions we did not observe a characteristic enrichment of other factors at hyper-accessible promoters but detected a pronounced enrichment of the heat shock factor 1 (Hsf1) and Spt7. Spt7 is a subunit of the SAGA complex, located upstream of the TSS upon heat shock, suggesting these promoters are kept accessible for rapid inducible activation (Figure S7D).

Next, we investigated the sub-nucleosomal fraction. We detected unstable nucleosomes not only at the TTS (Figure S8A), as already described (Chereji *et al*, 2017; Xi *et al*, 2011; Weiner *et al*, 2010), but also directly upstream of the TSS, like in *D. melanogaster* (Figure 7A, Supplemental Dataset 2). Next, we subdivided promoters of expressed genes into promoters with nucleosome-depleted regions (NDR) and promoters harboring an unstable nucleosome. In total 2,942 (53%) promoters are occupied by an unstable nucleosome, illustrating the abundance of this promoter type (Figure 7A). Our classification allowed us to detect sub-nucleosomal structures in independent H2B- and H3-MNase-ChIP data sets, even though higher MNase digestion conditions were used (Figure S8B) (Rossi *et al*, 2021). Those sub-nucleosomal patterns were absent using MNase-ChIP directed against histone modifications, such as H3K4me3, confirming that they are not a result of cross-linking artifacts. Remarkably, histone variant H2A.Z is enriched at unstable nucleosome promoters, agreeing with recent studies showing sub-nucleosomal structures in *S. cerevisiae* (Figure S8B) (Brahma & Henikoff, 2019; Xi *et al*, 2011). It was suggested that unstable nucleosomes are preferentially found at wide NDRs (> 150 bp distance between +1 and -1 nucleosome) (Kubik *et al*, 2015). Accordingly, we checked whether sub-nucleosomal fragments are preferentially enriched at wide NDRs but could not observe a significant difference (Figure S8C). As the main difference between the study of Kubik *et al*. and ours was that we used MNase-ChIP-seq data instead of sole MNase digestions, we reasoned that non-histone proteins protecting DNA accessibility might have biased their analysis. Indeed, the mono-nucleosome-sized fragments at wide NDRs accumulated an additional peak upstream of the TSS, which was not present in the H4-MNase-ChIP seq data (Figure S8D). Furthermore, the sub-nucleosome fraction revealed a high abundance of non-histone proteins upstream of the TSS, likely resulting in nucleosome-sized fragments at wide NDRs under low MNase conditions.

**Figure 7.**
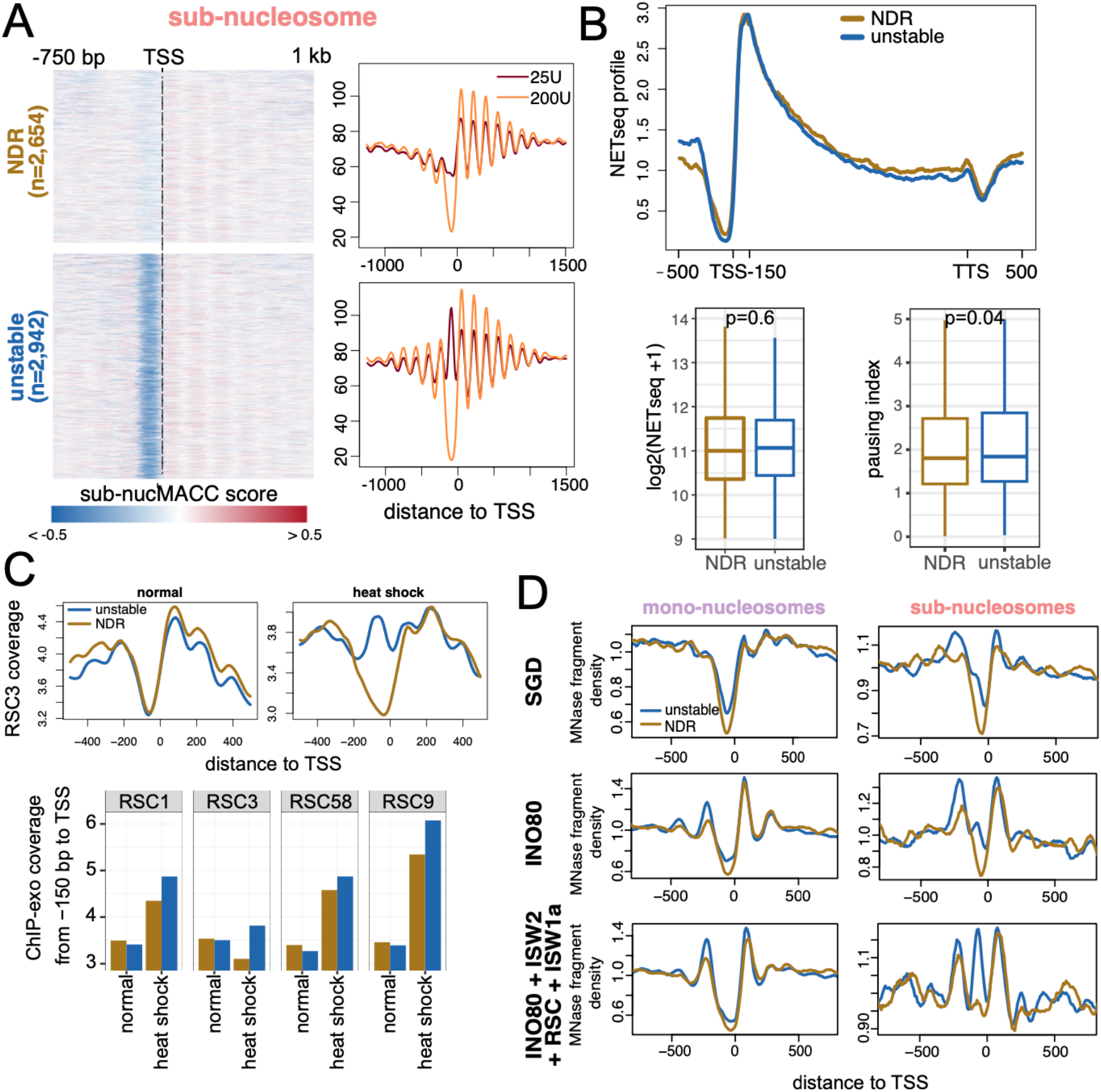
(A) Grouping of promoter types into promoter harboring an unstable nucleosome (unstable; n=2,942) or a nucleosome depleted region (NDR; n=2,654) in *S. cerevisiae*. Left panel: Heatmap showing the distribution of sub-nuccMACC scores. Right panel: Average plot showing the sub-nucleosomal fragment frequencies of the different MNase conditions. (B) Gene expression analysis of the promoter subgroups (unstable in blue and NDR in gold) using NETseq data. Top panel: Median NETseq profile showing nascent RNA abundance. Regions upstream of TSS, downstream of TTS and the first 150 bp downstream of TSS are unscaled. The remaining gene body is scaled to have the same length for each gene. Bottom panel: Boxplots showing the RNA Pol II initiation rate as calculated from the median NETseq signal over the +1 nucleosome (first 150 bp downstream of TSS) is shown on the left. The pausing index is shown on the right. Pausing index was calculated as the fold change of the NETseq signal in the first 150 bp downstream of the TSS versus the remaining gene body. (C) Occupancy analysis of RSC components at unstable nucleosome or NDR promoter types in the context of heat shock using ChIP-exo data (Rossi *et al*, 2021). Top panel shows the median normalized ChIP-exo coverage of the subunit RSC3 under normal and heat shock conditions. Bottom panel shows the median normalized ChIP-exo signal at the first 150 bp downstream of the TSS for RSC subunits RSC1, RSC3, RSC58, RSC9. (D) MNase fragment density of *in vitro* reconstituted nucleosome arrays at unstable nucleosomes or NDR promoter. MNase fragments were size selected into mono-nucleosomal (140 bp – 200 bp; left panel) and sub-nucleosomal fragments (< 140 bp; right panel). Nucleosomes were assembled onto DNA by salt gradient dialysis (SGD) and purified remodeler were added as indicated (Krietenstein *et al*, 2016).

The transcription initiation rate at NDR and unstable nucleosome promoters did not significantly differ (Figure 7B). Nevertheless, the pausing index was increased at the promoters harboring an unstable nucleosome as already seen in *D. melanogaster*.

Since it was suggested that RSC (remodeling the structure of chromatin) nucleosome remodeling leads to MNase-sensitive structures through partially unwrapped nucleosome intermediates (Brahma & Henikoff, 2019), we tested whether subunits of the RSC complex are overrepresented at the sites of unstable nucleosome promoters compared to NDR promoters using the ChIP-exo datasets (Rossi *et al*, 2021). We did not detect differences at normal conditions, however, upon heat-shock induced stress we identified several RSC subunits, being enriched at the sites of the unstable nucleosomes (Figure 7C).

Next, we addressed, whether chromatin remodelers are sufficient to generate unstable nucleosomes. Hence, we used MNase-seq data of genome-wide *in vitro* reconstitutions, where the activity of purified remodelers was tested (Krietenstein *et al*, 2016).

The nucleosomes, assembled by salt gradient dialysis (SGD) onto genomic DNA, exhibited a very low level of sub-nucleosomal structures at the unstable nucleosome promoters (Figure 7D). The number of sub-nucleosomal fragments increased upon the addition of INO80, creating wider NDRs by positioning the +1 and -1 nucleosomes. The addition of ISW2, RSC and ISW1 further increased the amount of sub-nucleosomal fragments, specifically at unstable nucleosome promoters. Such a distinction of sub-nucleosome fragments into two groups could not be achieved using the NDR width classification after Kubik *et al*. (Figure S8E). Nevertheless, INO80 alone seems to be sufficient to adjust the NDR width into either narrow or wide NDRs.

In summary, the nucMACC pipeline uncovered unstable nucleosomes upstream of the TSS of specific genes in *S. cerevisiae*. Unstable nucleosomes are formed by the activity of chromatin remodeling enzymes in reconstituted nucleosome arrays, suggesting that they generate partially unwrapped nucleosomal intermediates, or sub-nucleosomal structures, like hexasomes.

## Discussion

We describe a complete MNase-seq analysis pipeline called nucMACC, which is designed to analyze genome-wide nucleosome and sub-nucleosome positions, nucleosome accessibility and stability using two or more MNase titration conditions. It provides a systematic approach to select nucleosome positions with exceptional properties and playing regulatory roles in the genome. The systematic detection of unstable nucleosomes unveils their regulatory role in RNA Pol II promoter-proximal pausing and transcription factor binding.

MNase-seq is not a new method, and studies to analyze chromatin accessibility have previously been performed (Mieczkowski *et al*, 2016; Chereji *et al*, 2019). However, the use of these pipelines remains challenging to implement, as they use specific annotation files, which are only available for a few organisms, or require preinstalled software packages, some of which are not maintained anymore or require subscriptions. The nucMACC pipeline is publicly available and utilizes the nextflow workflow management system and Docker software containerization guaranteeing a seamless start on different computer systems and reproducible results (DI Tommaso *et al*, 2017). In addition, the nucMACC pipeline uncovers additional properties of structurally altered nucleosomes and sub-nucleosomes, resulting in new insights into chromatin function.

The nucMACC pipeline relates to the original MACC concept, which gives a qualitative and quantitative description of chromatin accessibility (Mieczkowski *et al*, 2016). We made two major modifications to obtain a higher resolution in mono-nucleosome analysis: 1) We selected only mono-nucleosomal fragments for the analysis by narrowing the fragment length selection to 140 – 200 bp. 2) Instead of subdividing the whole genome into arbitrary bins, we used the called nucleosome positions to measure chromatin accessibility. Consequently, every nucleosome is further characterized by its linker DNA accessibility. Nevertheless, the main findings of the original MACC study about chromatin accessibility could be confirmed with our pipeline.

A recent study used six different MNase titrations and the addition of spike-ins to accurately measure the kinetics of nucleosome release from chromatin (q-MNase-seq) (Chereji *et al*, 2019). The high number of MNase experiments and high sequencing depth requirements, make the data generation extremely laborious and costly, especially for organisms with large genomes. Here we established the minimal parameters to reveal chromatin accessibility and could show that the nucMACC pipeline works robustly with only two MNase titration conditions that should differ in about one order of MNase units applied. The use of spike-ins did not significantly improve the analysis and is therefore not required as the pipeline measures the relative change between the conditions. Nevertheless, to quantitively measure absolute nucleosome occupancy between various conditions spike-ins are compulsory. We could not compare the results of our nucMACC analysis directly with the q-MNase analysis (Chereji *et al*, 2019), as analyzed data tracks are not available, and we were not able to reproduce the results shown in the publication based on the provided method description. However, our analysis confirmed the results of the q-MNase analysis, that hetero-and euchromatic regions exhibit similar accessibility rates in *D. melanogaster* (Chereji *et al*, 2019), which has been initially observed in human cells (Schwartz *et al*, 2018). As a unique feature, nucleosomes with extraordinary chromatin accessibility are automatically detected in the nucMACC pipeline. Hyper-accessible nucleosomes are enriched at the promoter and correlate with DNase-seq assays, transcriptional activity, and active histone marks, whereas hypo-accessible nucleosomes show opposite characteristics.

So far MNase sensitive nucleosomes have been identified by direct comparison of high and low MNase conditions at a limited subset of selected genomic regions, such as TSS, Enhancers or CTCF binding sites (Ishii *et al*, 2015; Schwartz *et al*, 2018; Brahma & Henikoff, 2019; Kubik *et al*, 2015; Voong *et al*, 2016; Iwafuchi-Doi *et al*, 2016; Jeffers & Lieb, 2017; Xi *et al*, 2011). The nucMACC pipeline provides the first systematic, unbiased and genome wide approach to detect unstable nucleosomes. In agreement with previous reports, we find them preferentially upstream of the TSS of active genes. However, in contrast, we do not observe a correlation between transcription strength or NDR width and the presence of unstable nucleosomes (Kubik *et al*, 2015; Brahma & Henikoff, 2019; Voong *et al*, 2016; Ishii *et al*, 2015; Michl-Holzinger *et al*, 2022). As the positions of the denominated MNase-sensitive nucleosomes overlap with binding sites of non-histone factors at TSS, enhancer or transcription factor binding sites, we clearly show that it is necessary to use MNase combined with histone-ChIP datasets. Otherwise, DNA protected by non-histone proteins will result in nucleosome-sized fragments, preferentially at low MNase conditions and bias the results. Here we show that promoters in D. melanogaster or S. cerevisiae can be classified into two distinct promoter nucleosome configurations, promoters occupied by an unstable nucleosome upstream of the TSS, and nucleosome depleted promoters. Although their associated genes have similar transcript abundance, these promoter types are functionally different. We observed that genes associated with unstable nucleosomes had higher RNA pol II pausing rates in both *D. melanogaster* and *S. cerevisiae*. In line with other studies, we find unstable nucleosomes being associated with specific genes that are transcriptionally activated in response to stimuli (Jeffers & Lieb, 2017; Xi *et al*, 2011). Importantly, unstable nucleosomes are found in the promoters of genes before the induction of environmental changes, suggesting that these genes are poised for rapid upregulation. Therefore, we hypothesize that nucleosome stability is encoded at gene promoters, contributing to the high transcriptional plasticity observed at these sites. We could confirm our hypothesis using the *in vitro* reconstituted nucleosome arrays on the yeast genome (Krietenstein *et al*, 2016). In these datasets, we clearly observe that the addition of chromatin remodeling enzymes, results in the formation of unstable nucleosomes at the same subset of promoters as *in vivo.* These findings highlight the different nucleosome configuration capacities of promoter sequences in recruiting remodeling activities, which shape gene function.

About the exact structure and composition of unstable nucleosomes we can only speculate. In vertebrates, it was shown that histone variants like H3.3 or H2A.Z contribute to nucleosome instability (Jin & Felsenfeld, 2007). Indeed, we observed a partial enrichment of H2A.Z in yeast unstable nucleosomes, but this correlation is not sufficient to account for the stability of most unstable nucleosomes. Furthermore, we used cross-linked nucleosome data datasets, which would overturn the weakened histone DNA interactions observed for histone variants (Xi *et al*, 2011).

We show that unstable nucleosomes are enriched in sub-nucleosomal fragments and are not limited to wide NDRs, but can also be found at narrow NDRs, where the space is too short to assemble full nucleosomes. These findings indicate the existence of smaller, non-canonical sub-nucleosomal structures, like hexasomes or hemisomes (Chong & Gan, 2023). However, here we face one of the main limitations in MNase-seq data – looking at population wide averages. We cannot exclude the possibility that different nucleosome array configurations exist in individual cells. Here it would be interesting to investigate unstable nucleosome promoters further, using single-molecule DNA sequencing approaches to measure nucleosome positions on individual chromatin fibres (Abdulhay *et al*, 2020).

Another possibility might be, that unstable nucleosomes represent sub-nucleosomal intermediate states of the ATP dependent nucleosome remodeling process. It was reported that histones accumulate upstream of the TSS at unstable nucleosome promoters, upon mutation of the FACT subunit SPT16, indicating that those sites undergo active remodeling to promote RNA Pol II transcription through the +1 nucleosome (Obermeyer *et al*, 2023; Michl-Holzinger *et al*, 2022). Furthermore, partially unwrapped nucleosome structures were proposed during INO80 or RSC remodeling resulting in sub-nucleosomal structures (Brahma & Henikoff, 2019; Hsieh *et al*, 2022). In line with these studies, we observed preferential enrichment of RSC components at unstable nucleosome positions upon heat shock and an increase in the number of unstable nucleosomes, by adding nucleosome remodeler to reconstituted nucleosome arrays.

In summary, we provide a publicly available pipeline for analyzing nucleosome positions and nucleosomal properties using MNase titrations, which for the first time provides a systematic framework to identify unstable nucleosomes.

## METHODS

### Public datasets used in this study

#### Drosophila melanogaster

Whole chromatin MNase-seq, MNase-ChIP-H3-seq, MNase-ChIP-H4-seq and RNA-seq (GSE78984), M1BP-ChIP-seq (GSE101554), GRO-seq (GSE58955), and MNase-seq with spike-ins in S2 (GSE128689) were downloaded from the European nucleotide archive (ENA) database. Histone modification ChIP-seq data were retrieved through modENCODE, with the following accession numbers: H3K27ac (296), H3K27me3 (298), H3K4me1 (304), H3K9me3 (313), H3K4me3 (3761). DNase-seq (GSE172753) data were retrieved from the GEO database.

#### Saccharomyces cerevisiae

Whole chromatin MNase-seq, MNase-ChIP-H4-seq (GSE83123), NET-seq (GSE25107), *in vitro* MNase-seq on SGD assembled nucleosomes (GSE72106), ChIP-exo data of Rsc3, Rsc1, Rsc58, Rsc9, Spt7, Hsf1, Bdf1, CTDSer5, Ssl1, Ssl2, Taf2, Taf3, Tfa1, Tfb1, Tfb2, Tfb3, Tfb4, Tfg1 (GSE147927) were downloaded from the European nucleotide archive (ENA) database.

### The nucMACC pipeline

All the steps from fastq files to annotate (sub-)nucMACC scores were integrated and automated within the nucMACC pipeline, which is available on GitHub at uschwartz/nucMACC. The version of the nucMACC pipeline used in this study was v.1.2. The nucMACC pipeline was executed using nextflow, an open-source workflow management system, and runs the software within stable Docker containers, ensuring a reproducible analysis workflow (DI Tommaso *et al*, 2017).

Raw sequencing data in the form of fastq files were first subjected to quality control checks using FastQC (Andrews, 2010). Reads were mapped against the reference genome (dm3 / sacCer3) using Bowtie2 with the following parameters --very-sensitive-local --no-discordant (Langmead & Salzberg, 2012). Alignments with mapping quality (MAPQ) below 30 were removed using samtools (Li *et al*, 2009). Next, the alignments were filtered based on their size to categorize them into mono-nucleosomes (140-200 bp) or sub-nucleosomes (< 140 bp) using the alignmentSieve function from the deepTools suite (Ramírez *et al*, 2016). Additionally, any blacklisted regions were removed during this step to ensure that only reliable regions were considered for further analysis. Nucleosome positions were identified using DANPOS2 (Chen *et al*, 2013). For this, pooled mono-nucleosome samples were used to call the positions of mono-nucleosomes, while sub-nucleosome positions were called from samples with the lowest MNase concentration. Here library size was normalized to the effective genome size, paired-end reads centered to their midpoint and extended to 70 bp to obtain nucleosome-sized positions. The GC content at the identified nucleosome positions was measured using bedtools genomecov function (Quinlan & Hall, 2010). To quantify the fragments associated with each nucleosome position, the featureCounts tool was employed (Liao *et al*, 2014). Nucleosome positions with low fragment counts were filtered out to ensure robustness in the downstream analysis. Specifically, mono-nucleosome positions with fragment count less than 30 and sub-nucleosome positions with counts less than 5 were removed.

### nucMACC score calculation

First the fragment counts were normalized based on the library size to obtain counts per million (CPM). A pseudocount was determined as the median of the sample median count. A linear regression analysis was performed between log2-transformed MNase concentration and the log2-transformed normalized counts with the added pseudocount. The raw nucMACC scores were estimated from the slope obtained from the regression analysis multiplied by minus one. Next, the raw nucMACC scores were corrected for the underlying GC content. Nucleosome positions exhibiting rarely occurring extreme GC content were filtered out. A locally weighted scatterplot smoothing (LOESS) fit was calculated for the filtered GC content and corresponding raw nucMACC scores using a smoothing parameter α of 0.1. Raw nucMACC scores were subtracted by the deviation of the loess fit from the median raw nucMACC score at the corresponding GC content. The sub-nucMACC scores were determined in the same way, except that the slope values were not multiplied by minus one to represent the stability scores.

### Identification of hypo-/hyper-accessible nucleosomes

To identify hypo- and hyper-accessible nucleosomes, the nucMACC scores were normalized to a deviation from the highest to lowest value of 1 and plotted against the ranks of the nucMACC scores, which were normalized by the total number of ranks. A LOESS smoothing was applied and the first derivate of the LOESS fit was calculated to deduce the slope of the curve. Hypo-accessible nucleosomes are determined as the lowest nucMACC scores until the slope of the curve falls below 1 and hyper-accessible nucleosomes are the highest nucMACC scores after the slope of the curve exceeds again a slope of 1.

### Identification of unstable and non-canonical nucleosomes

Unstable and non-canonical nucleosomes were similarly determined as hyper-/hypo-accessible nucleosomes, except that sub-nucleosome positions are selected, which either are unique or exhibit at least four-fold more normalized counts in the lowest MNase condition compared to the mono-nucleosomes. Unstable nucleosomes are determined as the lowest sub-nucMACC scores until the slope of the curve falls below 1 and non-canonical nucleosomes are the highest sub-nucMACC scores after the slope of the curve exceeds again a slope of 1.

### Spike-in analysis

We compared the results of the nucMACC pipeline ignoring the spike-in information with a slightly modified version using the spike-ins for normalization. Therefore, reads were aligned to the sacCer3 *Saccharomyces cerevisiae* genome and treated analogously to the nucMACC pipeline. The number of fragments per mono- and sub-nucleosome fraction was counted, and normalization factors for the nucMACC analysis in *D.melanogaster* were calculated by dividing the number of *S. cerevisiae* derived fragments by 1 Mio.

### Downstream analysis

The code used to generate the figures in this manuscript can be found in the project GitHub repository uschwartz/nucMACC_Paper. Density correlation scatterplots were generated using the LSD R package (Schwalb *et al*, 2020). Heatmaps and metaplots were calculated using the computeMatrix, plotHeatmap, plotProfile functions of the deeptools suit (Ramírez *et al*, 2016) and further modified in R using basic plot functions or the pheatmap package (Kolde, 2019). Genomic features were annotated using the Bioconductor package ChIPseeker (Yu *et al*, 2015). Motif enrichment analysis was carried out using the findMotifs.pl script from the HOMER software suite (Heinz *et al*, 2010). Motifs from the integrated fly database were tested for enrichment at the TSS +/-100 bp of genes associated either with an unstable nucleosome or a nucleosome depleted region. The DNA sequence around the TSS of all expressed genes served as background.

Gene set over-representation analysis was carried out using the Metascape online tool (Zhou *et al*, 2019). Enrichment was performed using the Gene Ontology (GO) database for biological processes (BP) and restricted to terms with at least 20 genes overlapping to the tested gene set.

### GRO-seq / NET-seq analysis

GRO- and NET-seq data have been analyzed using the in-house nextflow transcriptomics pipeline available at GitHub (uschwartz/RNAseq).

Initially, quality control of the raw sequence reads was conducted using FastQC (v0.11.8) (Andrews, 2010). Subsequently, reads were mapped to the reference genome and corresponding gene annotation using the Spliced Transcripts Alignment to a Reference (STAR) software (version v2.7.8a)(Dobin *et al*, 2013). The following options were used to optimize the alignment process: --outFilterType BySJout, --outFilterMultimapNmax 20, -- alignSJoverhangMin 8, --alignSJDBoverhangMin 1, --outFilterMismatchNmax 999, -- alignIntronMin 10, --alignIntronMax 1000000, --outFilterMismatchNoverReadLmax 0.04, -- runThreadN 12, --outSAMtype BAM SortedByCoordinate, --outSAMmultNmax 1, and -- outMultimapperOrder Random.

Post-mapping quality control was performed using the rnaseq analysis mode of Qualimap (v2.2.1)(García-Alcalde *et al*, 2012). The level of PCR duplication was assessed using Picard MarkDuplicates (v2.21.8) (Broad Institute, 2019), and dupRadar (version v1.15.0) (Sayols *et al*, 2016). Gene expression quantification was carried out using featureCounts (v1.6.3) (Liao *et al*, 2014).

### ChIP-exo data processing

ChIP-exo sequencing data in *S. cerevisiae* was downloaded and trimmed with TrimGalore v0.6.7 (github.com/FelixKrueger/TrimGalore). Trimmed reads were aligned to the sacCer3 reference genome using bowtie2 v2.4.2 (Langmead & Salzberg, 2012) with the following options “--very-sensitive -I 0 -X 500 --fr --no-mixed --no-discordant –dovetail”. Duplicate reads were identified and removed using picard v2.18.7 (Broad Institute, 2019). Bigwigs mimicking standard ChIP were generated using the bamCoverage function from deeptools v3.5.1 (Ramírez *et al*, 2016) with the options: “-bs 10 --normalizeUsing CPM --extendReads -- centerReads --minMappingQuality 1”.

## ACKNOWLEDGEMENTS

The study was financed by the DFG SFB960 (to GL and US). We thank Philipp Korber for fruitful discussions and Joachim Griesenbeck for helpful comments on this manuscript.

## AUTHOR CONTRIBUTIONS

***Wernig-Zorc Sara:*** Methodology, Validation, Formal analysis, Investigation, Visualization, Writing – Review & Editing ***Kugler Fabian:*** Methodology, Software, Validation, Formal analysis, Investigation, Visualization ***Schmutterer Leo:*** Software, Formal analysis, Investigation ***Räß Patrick:*** Software ***Hausmann Clemens:*** Software ***Holzinger Simon:*** Methodology, Formal analysis, Writing – Review & Editing ***Längst Gernot:*** Conceptualization, Funding Acquisition, Resources, Project administration, Supervision, Writing – Review & Editing ***Schwartz Uwe:*** Conceptualization, Methodology, Software, Validation, Formal analysis, Investigation, Writing – Original Draft, Supervision, Visualization

