## SupplementaryData for "nucMACC: A MNase-seq pipeline to identify structurally altered nucleosomes in the genome"

### Supplemental Data of:
### nucMACC: An optimized MNase-seq pipeline measures genome-wide nucleosome accessibility and stability

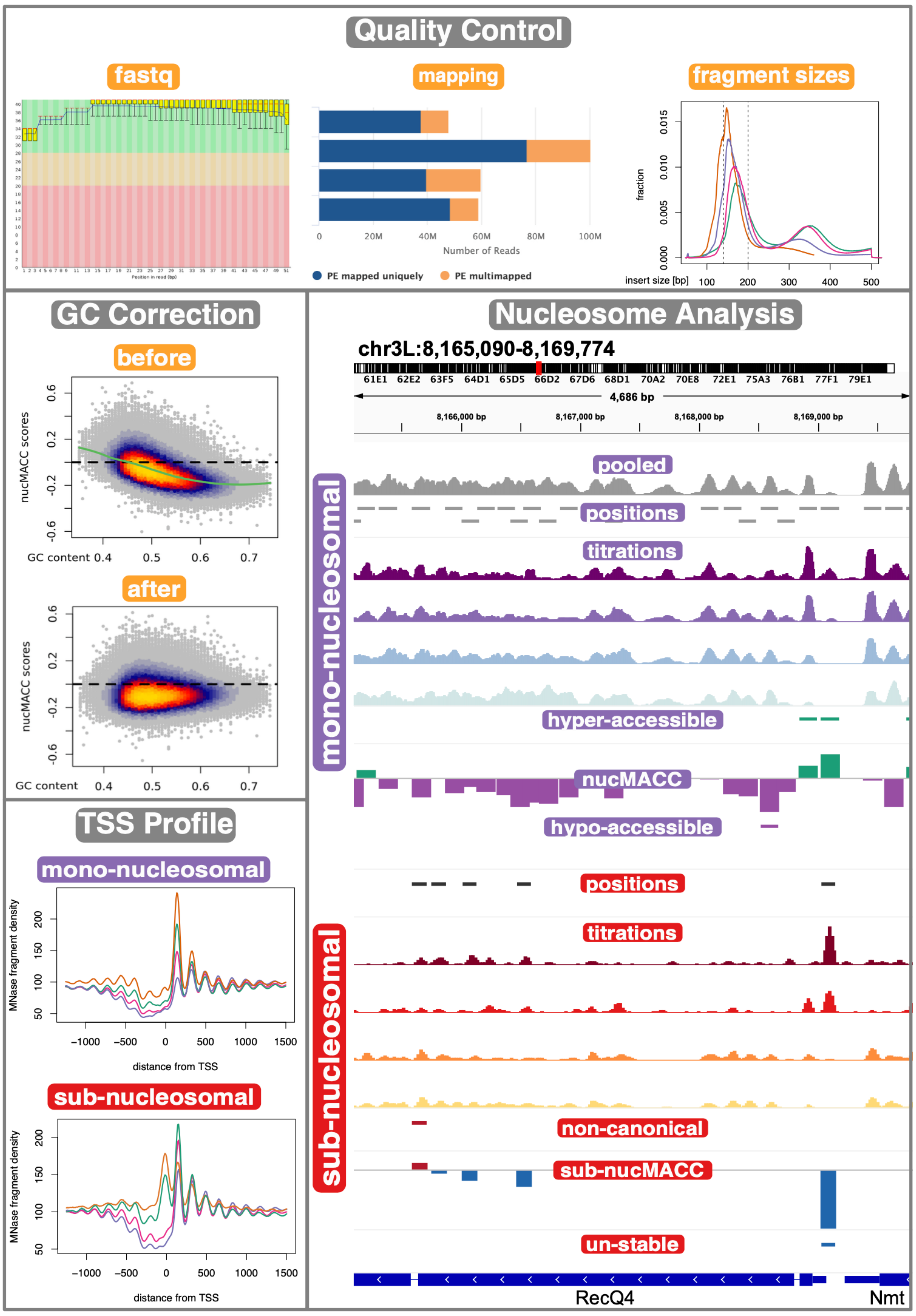


**Figure S1**

Overview of nucMACC pipeline. Quality reports are generated at different steps of the pipeline to control each step of the analysis and the underlying data (top panel). The pipeline generates several output files, such as nucleosome positions, MNase profiles, (sub-)nucMACC scores, or special nucleosome features, which can be directly visualized in the genome browser (right panel). During the calculation of (sub-)nucMACC scores, the GC bias of MNase is corrected based on a LOESS fit (green line) (middle left panel). Optional fragment profiles at the TSS are generated (bottom left panel).

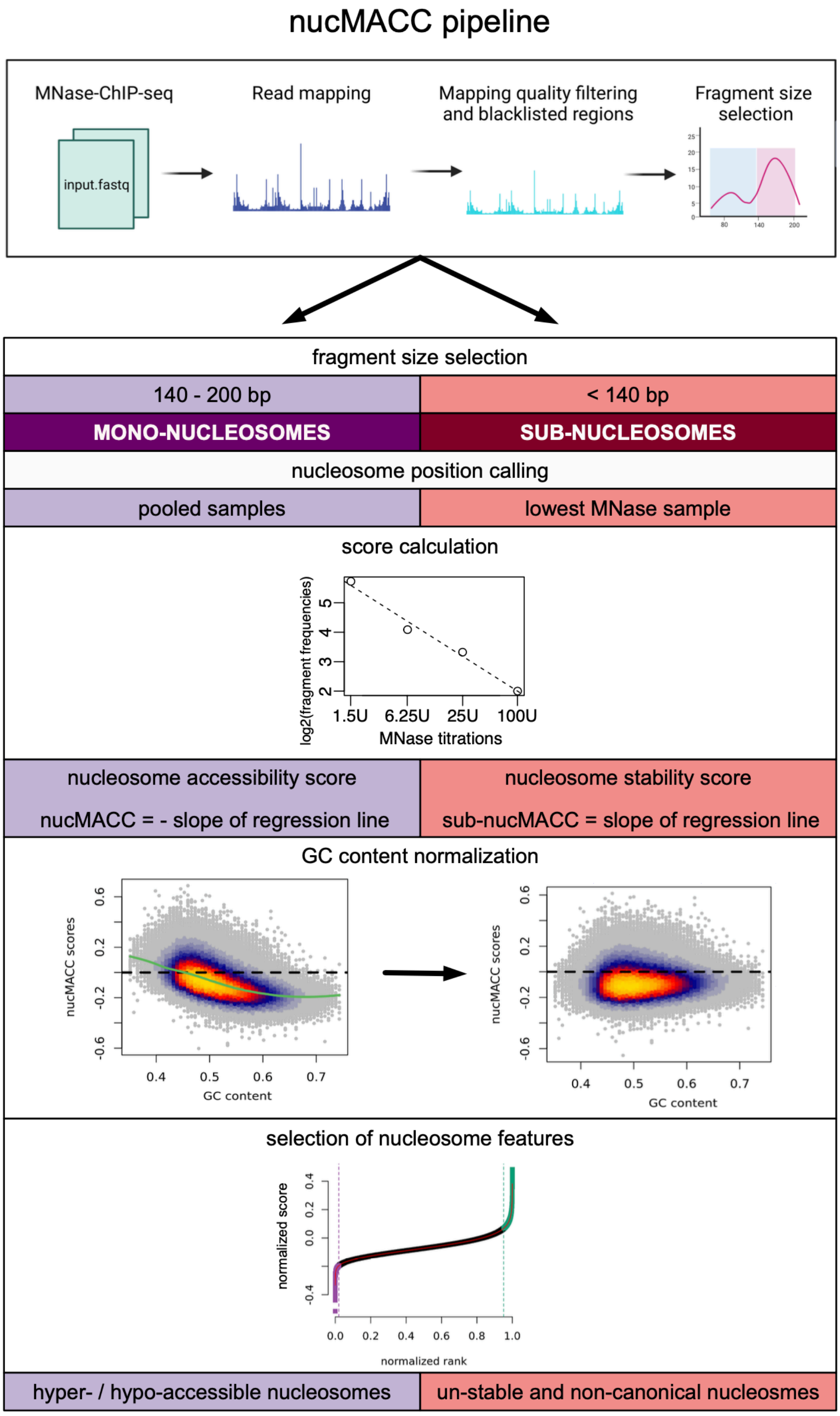


**Figure S2**

Workflow of the nucMACC pipeline. The nucMACC pipeline starts with raw data in .fastq format and requires a minimum of two MNase conditions. Fragments are mapped to the reference genome and aligned fragments are filtered based on mapping quality. In the next step, fragments are divided into sub-nucleosomal and mono-nucleosomal-sized fragments, which are processed separately. For nucleosome position calling all MNase concentration conditions are pooled, whereas for sub-nucleosomes, only the lowest MNase titration is used to call sub-nucleosomal positions. Sub-nucleosomal positions are consequently filtered by previously called mono-nucleosomal positions to obtain enriched sub-nucleosome positions. Then a nucMACC score is calculated by counting fragments per each nucleosome position and each MNase condition. The slope of a linear regression fit is determined and represents the raw score, which is normalized to the underlying GC% content in the following step. The normalized score is referred to as the nucleosome MNase accessibility score (nucMACC) or sub-nucMACC score, respectively. In the final step, special nucleosome groups are obtained by analyzing where the (sub-)nucMACC score considerably deviates from the mean. From the mono-nucleosome fraction we obtain stable, hypo-, and hyper-accessible nucleosomes. While the special nucleosomes from the sub-nucleosomal fraction represent un-stable and non-canonical nucleosomes.

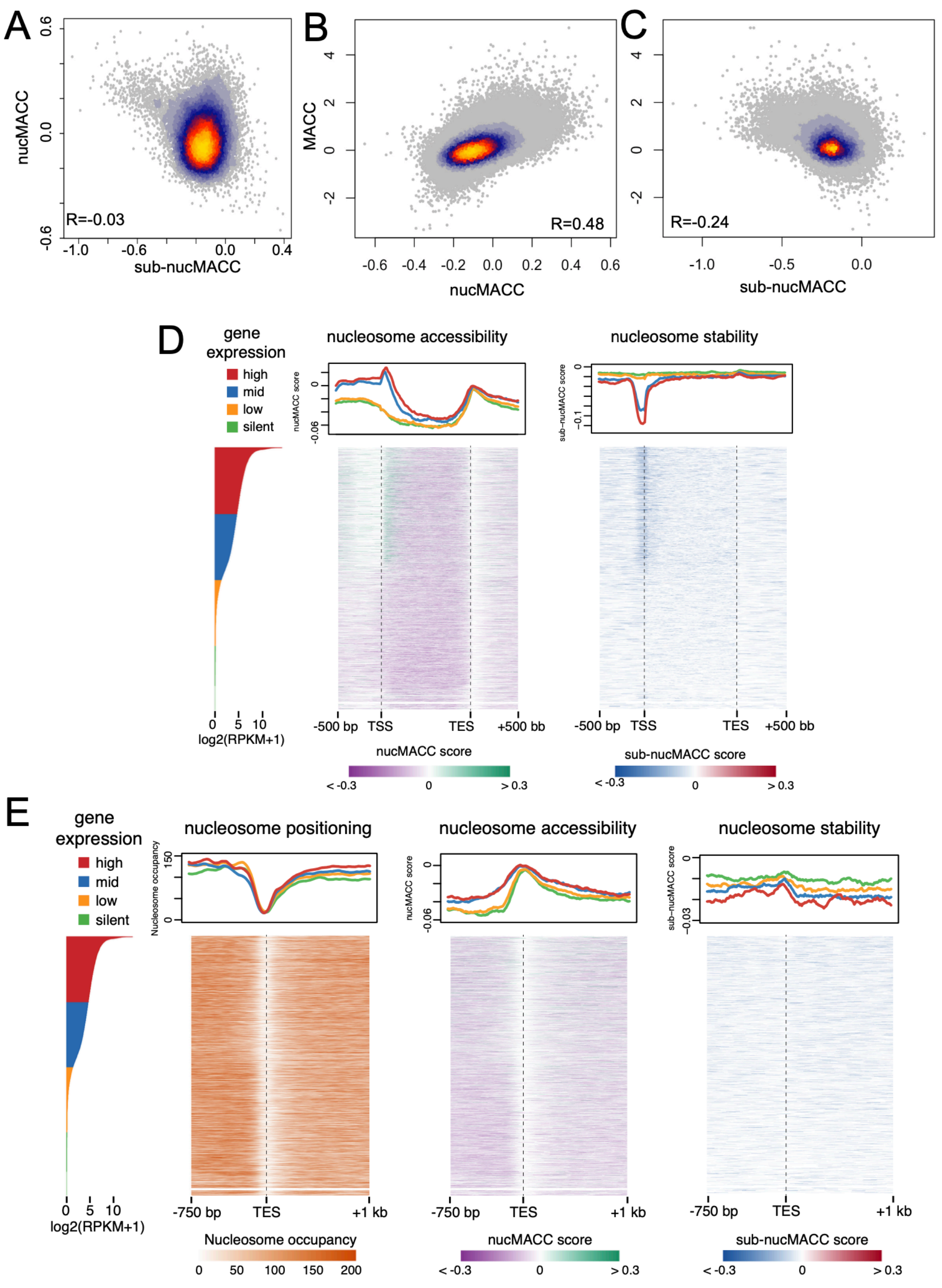


**Figure S3**

(A-C) Density Scatterplot showing correlation between (A) sub-nucMACC and nucMACC scores, (B-C) original MACC score after (Mieczkowski *et al*, 2016) and either (B) nucMACC or (C) sub-nucMACC scores. Pearson correlation coefficients (R) are indicated. (D-E) Heatmaps sorted by gene expression showing nucleosome accessibility (nucMACC score), and stability (sub-nucMACC score) over scaled gene bodies (D) or at the transcription end site (TES) including nucleosome positioning (E). Genes were subdivided by gene expression quartiles, as indicated by the colors.

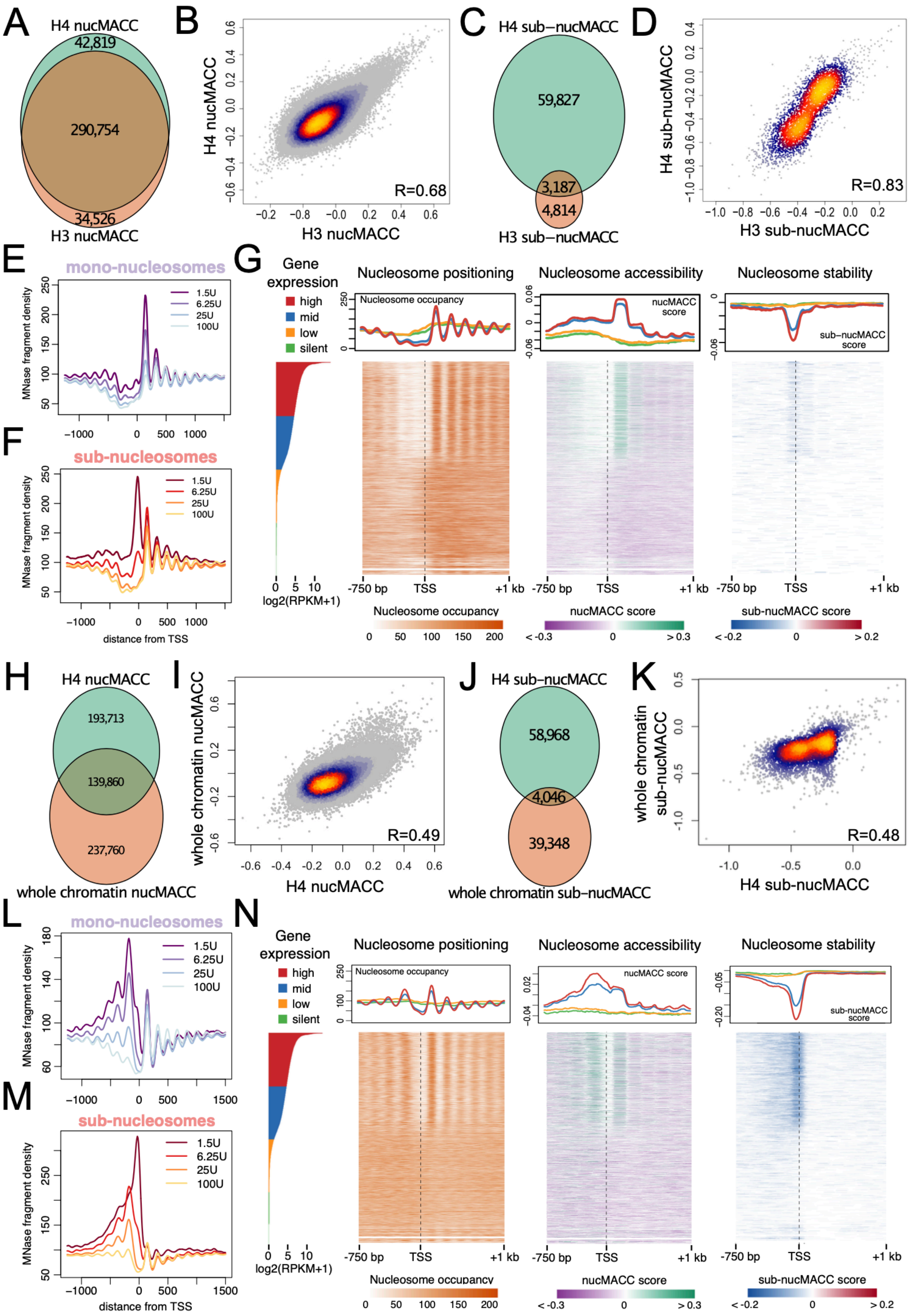

A
H4 nucMACC
42,819
290,754
34,526
H3 nucMACC
B
H4 nucMACC
H3 nucMACC
R=0.68
C
H4 sub−nucMACC
59,827
3,187
4,814
H3 sub−nucMACC
D
H4 sub−nucMACC
H3 sub−nucMACC
R=0.83
E
mono-nucleosomes
MNase fragment density
1.5U
6.25U
25U
100U
F
sub-nucleosomes
distance from TSS
G
Gene expression
high
mid
low
silent
log2(RPKM+1)
Nucleosome positioning
Nucleosome occupancy
Nucleosome accessibility
nucMACC score
Nucleosome stability
sub−nucMACC score
-750 bp
TSS
+1 kb
H
H4 nucMACC
193,713
139,860
237,760
whole chromatin nucMACC
I
whole chromatin nucMACC
H4 nucMACC
R=0.49
J
H4 sub−nucMACC
58,968
4,046
39,348
whole chromatin sub−nucMACC
K
whole chromatin sub−nucMACC
H4 sub-nucMACC
R=0.48
L
mono-nucleosomes
M
sub-nucleosomes
distance from TSS
N
Gene expression
Nucleosome positioning
Nucleosome accessibility
Nucleosome stability

**Figure S4**

(A) and (C) Venn diagrams showing the overlap of mono-nucleosome (A) or sub-nucleosome (C) positions determined using MNase combined with either H3 or H4 immunoprecipitations experiments in *D. melanogaster*. (B) and (D) Density scatterplot showing correlation of nucMACC (B) or sub-nucMACC (D) scores at overlapping positions between MNase combined with either H3 or H4 immunoprecipitations. (E-F) Average fragment frequencies at the TSS of MNase H3 data sets. (G) Heatmaps sorted by gene expression showing nucleosome positioning (left), accessibility (middle, nucMACC scores), and stability (right, sub-nucMACC scores) at TSS of MNase H3 data sets. (H) and (J) Venn diagrams showing the overlap of mono-nucleosome (H) or sub-nucleosome (J) positions determined using MNase either with H4 immunoprecipitation or without additional histone immunoprecipitation step (whole chromatin). (I) and (K) Density scatterplot showing Pearson correlation of nucMACC (I) or sub-nucMACC (K) scores at overlapping positions between MNase either with H4 immunoprecipitation or whole chromatin. (L-M) Average fragment frequencies at the TSS of MNase whole chromatin data sets. (N) Heatmaps sorted by gene expression showing nucleosome positioning (left), accessibility (middle, nucMACC scores), and stability (right, sub-nucMACC scores) at TSS of MNase whole chromatin data sets.

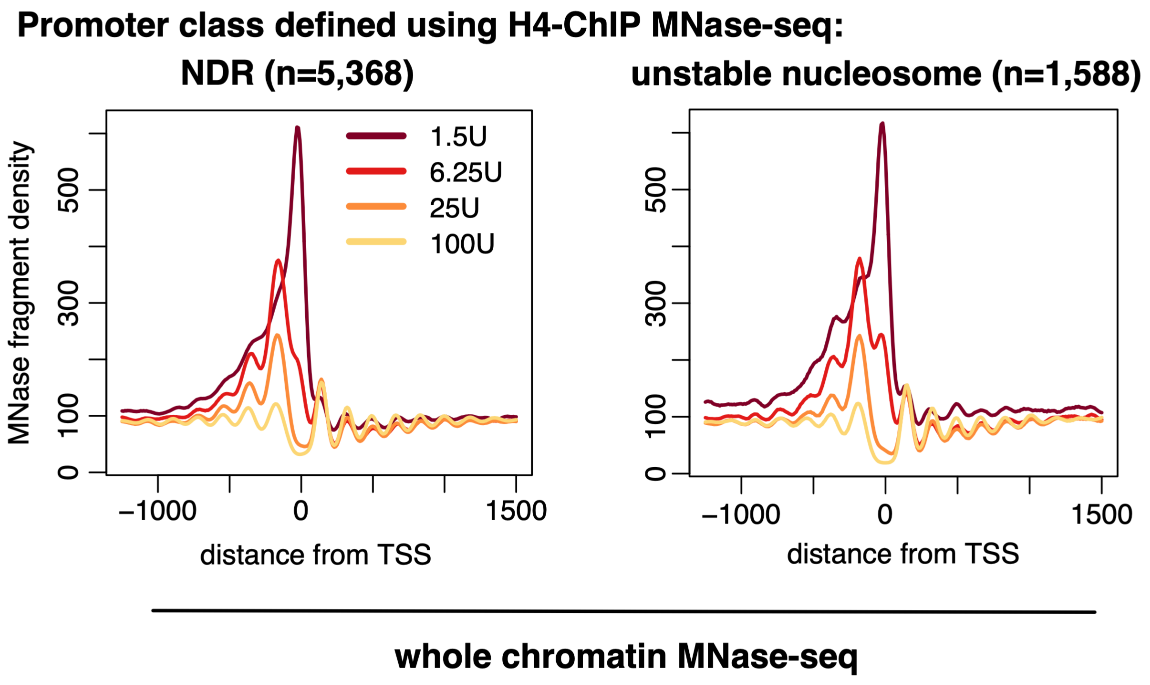


**Figure S5**

Average sub-nucleosomal fragment frequencies of MNase whole chromatin data sets at the TSS of expressed genes either exhibiting a nucleosome depleted region (NDR) (n=5,368, left) or containing an unstable nucleosome (n=1,588, right) directly upstream of the TSS.

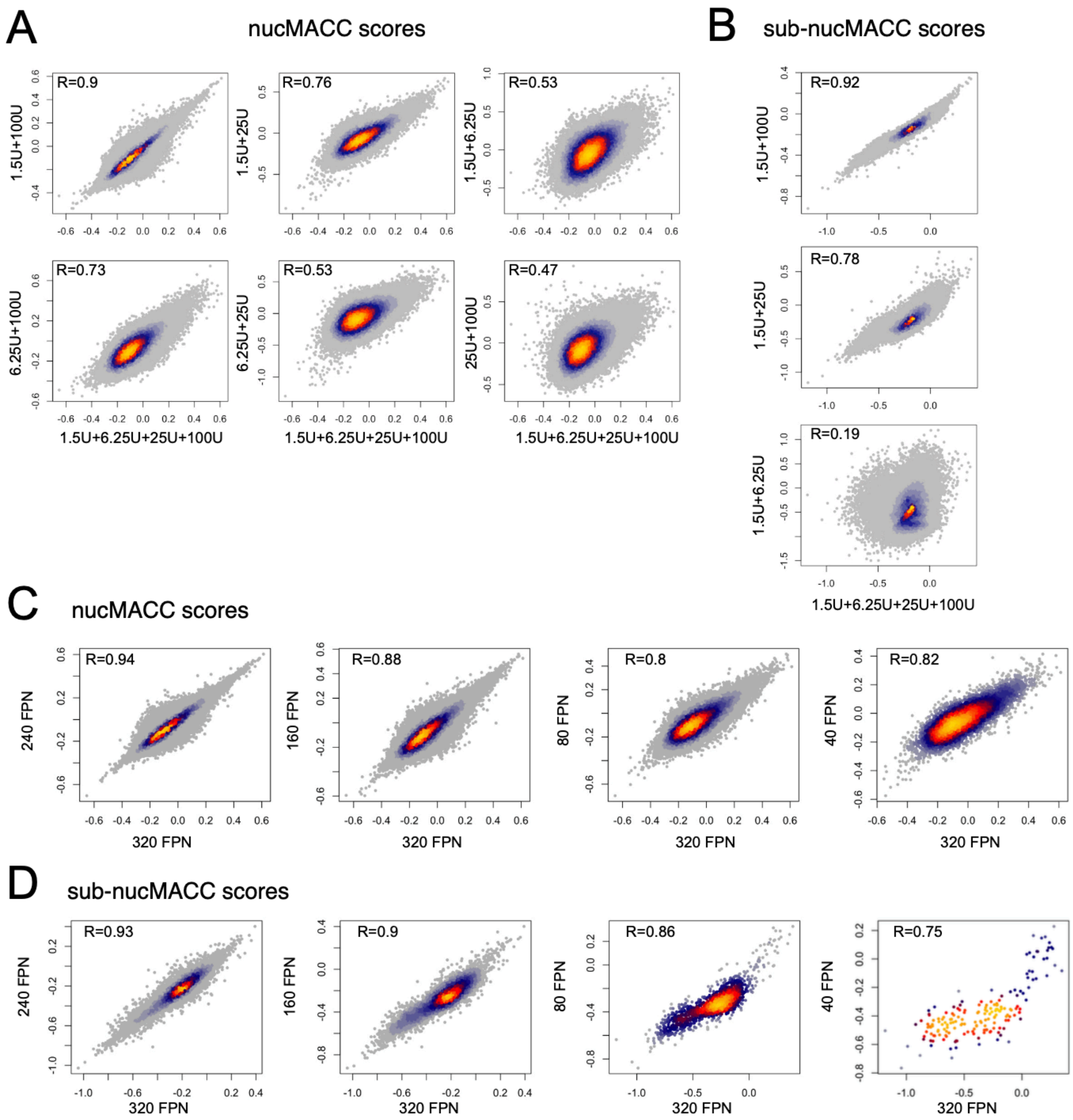


**Figure S6**

(A-B) Density scatterplots showing correlations of nucMACC (A) or sub-nucMACC (B) scores between all titrations and selected MNase titration pairs as indicated. (C-D) Density scatterplots showing correlations of nucMACC (C) or sub-nucMACC (D) scores between the total number of sequenced fragments (320 FPN) and subsampled fragments as indicated. FPN refers to the sequencing depth calculated as the number of fragments per nucleosome.
The Pearson correlation coefficient is indicated at the top.

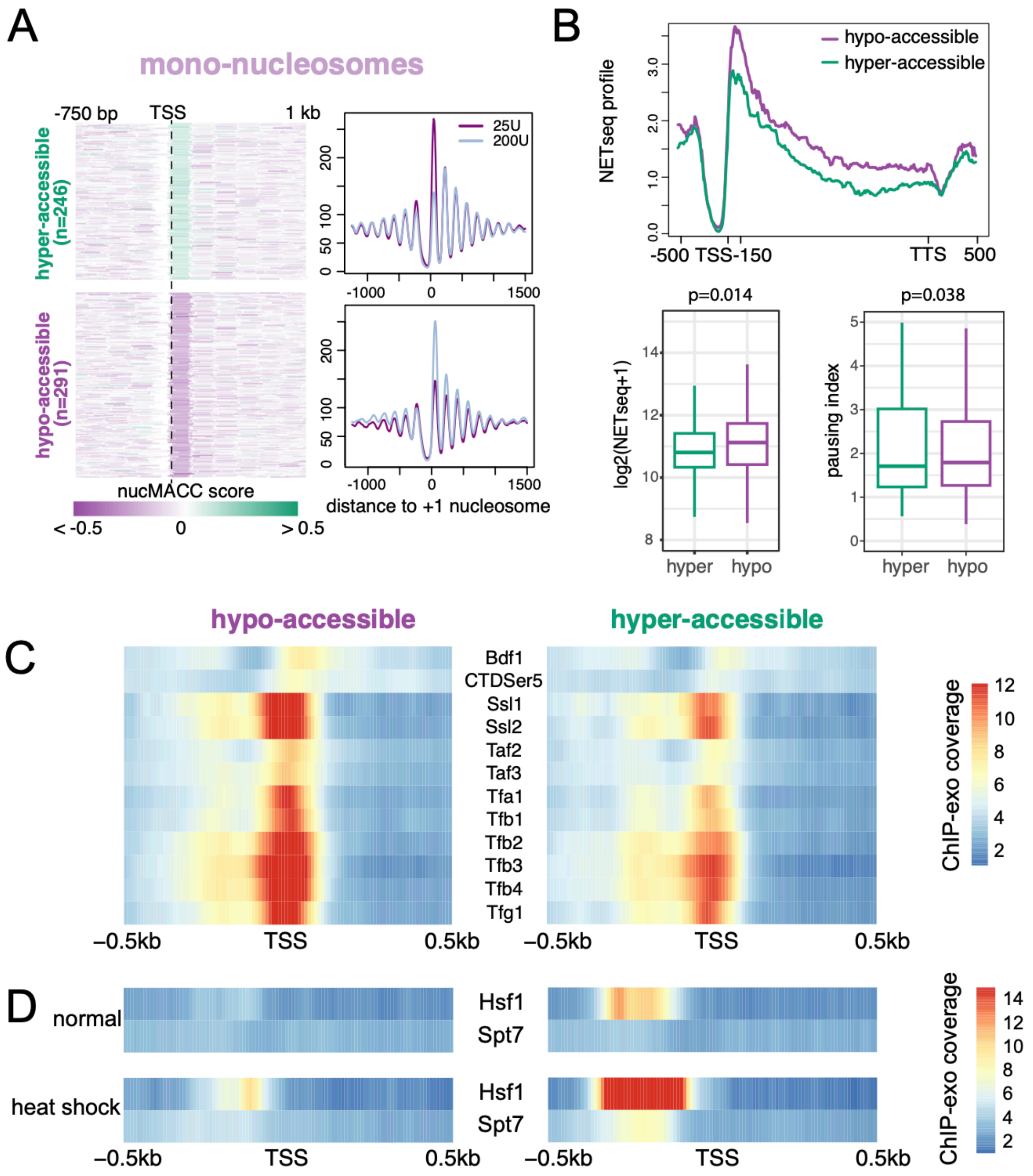


**Figure S7**

(A) Grouping of promoter types into promoter harboring a hyper- (n=246) or a hypo-accessible (n=291) +1 nucleosome in *S. cerevisiae*. Left panel: Heatmap showing the distribution of nuccMACC scores. Right panel: Average plot showing the mono-nucleosomal fragment frequencies of the different MNase conditions. (B) Gene expression analysis of the +1 nucleosome subgroups (hyper- in green and hypo-accessible in purple) using NETseq data. Top panel: Median NETseq profile showing nascent RNA abundance. Regions upstream of TSS, downstream of TTS and the first 150 bp downstream of TSS are unscaled. The remaining gene body is scaled to have the same length for each gene. Bottom panel: Boxplots showing the RNA Pol II initiation rate as calculated from the median NETseq signal over the +1 nucleosome (first 150 bp downstream of TSS) is shown on the left. The pausing index is shown on the right. Pausing index was calculated as the fold change of the NETseq signal in the first 150 bp downstream of the TSS versus the remaining gene body. (C-D) Heatmap illustrating occupancy of certain factors at hyper-/hypo-accessible +1 genes around the TSS. The color scale indicates the normalized coverage in ChIP-exo experiments (Rossi *et al*, 2021).

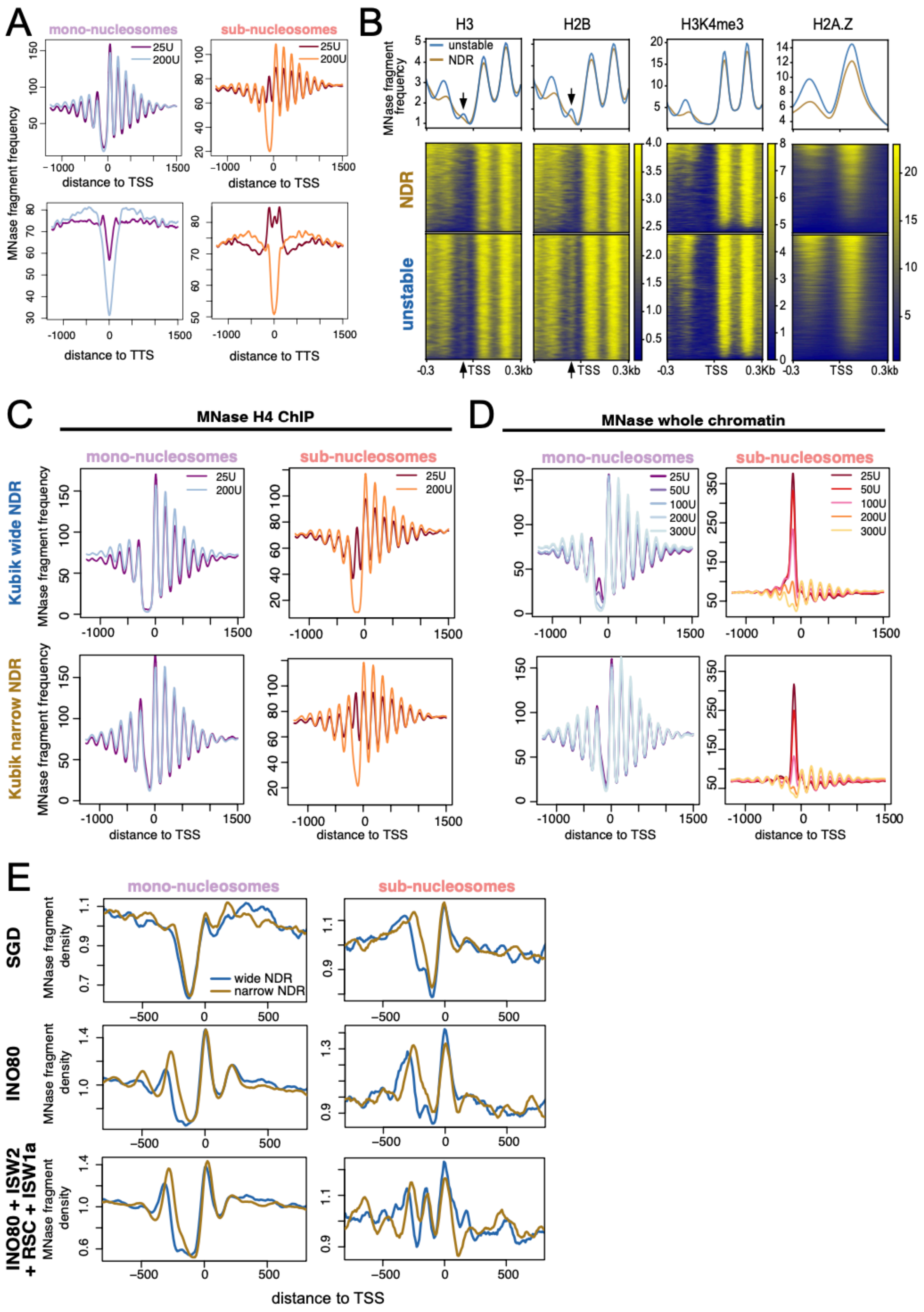

A
mono-nucleosomes
sub-nucleosomes
25U
200U
MNase fragment frequency
distance to TSS
distance to TTS
B
H3
H2B
H3K4me3
H2A.Z
unstable
NDR
MNase fragment frequency
TSS
C
MNase H4 ChIP
Kubik wide NDR
Kubik narrow NDR
D
MNase whole chromatin
50U
100U
300U
E
SGD
INO80
INO80 + ISW2 + RSC + ISW1a
MNase fragment density
wide NDR
narrow NDR
distance to TSS


**Figure S8**

(A) Average fragment frequencies of mono- (left) and sub-nucleosomal (right) fragments at the TSS (top) or TTS (bottom) in *S.cerevisiae*. (B) Meta-plot and heatmap showing fragment abundance at NDR and unstable nucleosome promoter. MNase-ChIP data sets without fragment size selection were used from (Rossi *et al*, 2021). Arrows indicate the expected position of unstable nucleosomes. (C-D) Average fragment frequencies of MNase-H4-ChIP (C) or MNase whole chromatin (D) data sets at the TSS subgrouped by NDR width after (Kubik *et al*, 2015). (E) MNase fragment density of *in vitro* reconstituted nucleosome arrays at unstable narrow or wide NDRs. MNase fragments were size selected into mono-nucleosomal (140 bp – 200 bp; left panel) and sub-nucleosomal fragments (< 140 bp; right panel). Nucleosomes were assembled onto DNA by salt gradient dialysis (SGD) and purified remodeler were added as indicated (Krietenstein *et al*, 2016).
